# Pre-dauer starvation rapidly and reversibly reduces niche proliferative signaling to the *C. elegans* germ line

**DOI:** 10.1101/2025.05.23.655783

**Authors:** Fred A. Koitz, Camille P. Miller, Brian Kinney, Kacy Lynn Gordon

## Abstract

Early life stresses impact reproductive outcomes in many organisms. In response to crowding and starvation, *C. elegans* nematodes form dauer larvae, in which development arrests until conditions improve. We discovered dramatic differences in gonad size and germ cell number among dauers that form under different conditions. We used live cell imaging of fluorescent proteins in otherwise wild-type and mutant animals combined with food-removal, recovery, and brood-size assays to investigate the causes and consequences of this germline difference. Pre-dauer feeding, but not nutrient sensing via the DAF-2/insulin-like signaling receptor or DAF-7/TGF-β, is required for plasticity in gonad size. Gonad differences in dauer have lifelong reproductive consequences; severely starved worms make small dauer gonads and have small broods. Pre-dauer starvation induces germline quiescence and near-instantaneous reduction of the Notch ligand LAG-2 on the germline stem cell niche. A rapid return to germline Notch dependence and an increase in presentation by the germline stem cell niche of LAG-2–independent of *lag-2* transcriptional upregulation–are among the earliest events of dauer recovery.

**SUMMARY STATEMENT:** Pre-dauer starvation arrests gonad growth, reduces presentation of the germline niche cue LAG-2, and diminishes future reproductive success. Dauer recovery rapidly restores LAG-2 ligand and germline Notch dependence.

## INTRODUCTION

Lifelong development requires environmental responsiveness within the developmental program. A special case of developmental response to difficult environmental conditions is the diapause state, in which development arrests in unfavorable conditions and resumes when conditions improve (Harsimran Kaur Gill et al., 2017; Renfree and Fenelon, 2017). Obligate diapause states are built into an animal’s life cycle in anticipation of poor conditions, while facultative diapause states respond to an organism’s environment (Hand et al., 2016; Wilsterman et al., 2021). One well-studied facultative diapause state is the *C. elegans* dauer (Cassada and Russell, 1975). Dauer larvae can withstand poor environmental conditions (Cassada and Russell, 1975; Golden and Riddle, 1984) for many times the typical lifespan of a worm (Klass and Hirsh, 1976). Improved environmental conditions trigger exit from dauer, resumption of development, and a (mostly) normal reproductive life (Cassada and Russell, 1975; Klass and Hirsh, 1976). Famously, time spent in the dauer larval stage does not negatively affect adult lifespan (Hirsh et al., 1976; Klass and Hirsh, 1976).

It is the preservation of fertility rather than increased total lifespan alone that makes dauer adaptive. Previous studies discovered different long-term effects on post-dauer reproduction depending on the dauer-inducing conditions worms experienced (Gimond et al., 2025; Hall et al., 2010; Kim and Paik, 2008; Klass and Hirsh, 1976; Ow et al., 2018; Webster et al., 2018), focused on recovery. We set out to investigate the cells of the gonad and germ line during the dauer larval period itself to look for developmental antecedents of later reproductive differences.

*C. elegans* goes through four larval stages (L1-L4) before a terminal molt into the reproductive adult (Corsi et al., 2015). The dauer is an alternate third larval stage. Entry into the dauer developmental pathway is specified in the L1 larval stage by low food and high pheromone, and is followed by L2d, which lasts longer than the standard L2 (Golden and Riddle, 1984). In persistently poor conditions, the L2d molts into a non-feeding dauer larva with a thick cuticle, arrested development including absence of germ cell proliferation, and altered metabolism (Cassada and Russell, 1975; Erkut and Kurzchalia, 2015; Golden and Riddle, 1984; Wadsworth and Riddle, 1989).

The *C. elegans* hermaphrodite gonad develops post-embryonically from a four-cell primordium of two somatic blast cells and two primordial germ cells (Kimble and Hirsh, 1979). In the L1, a distal tip cell (DTC) is born from each of the somatic lineages and the primordial germ cells begin to divide (Kimble and Hirsh, 1979). The germ line relies on DTC expression of LAG-2, a Delta/Serrate/LAG-2 (DSL) ligand of the Notch signaling pathway, to maintain distal germ cells in an undifferentiated state, making the DTC the germline stem cell niche (Kimble and Crittenden, 2005). The developmental arrest of gonad and germ cell lineages in the dauer larva have been investigated genetically (Narbonne and Roy, 2006; Tenen and Greenwald, 2019), primarily using genetic mutants that constitutively form dauers at high temperature, called daf-c mutants.

Among these are mutations to genes in the insulin-IGF-1 signaling (IIS) pathway, including the gene encoding the sole *C. elegans* IIS receptor *daf-2* (Kimura et al., 1997). IIS integrates nutritional status with metabolism, developmental control, reproduction, and aging in *C. elegans* (Ewald et al., 2018; Murphy and Hu, 2013) and in other organisms, including humans (Barbieri et al., 2003; Claeys et al., 2002; Das and Arur, 2017). A “class 2” reduction-of-function allele, *daf-2(e1370),* is experimentally used to induce dauer larvae at high temperature (Gems et al., 1998; Karp, 2018; Kenyon et al., 1993). Another reduction-of-function allele, *daf-7(e1372)*/TGF-β, affecting a parallel pathway leading to dauer entry, also causes constitutive dauer formation at high temperature (Karp, 2018; Pierce et al., 2001; Ren et al., 1996). The TGF-β pathway links external cues to the reproductive system (Dalfó et al., 2012; Park et al., 2010, 2021; Pekar et al., 2017).

The germ line is especially sensitive to starvation throughout life (Angelo and Van Gilst, 2009; Dalfó et al., 2012; Pekar et al., 2017; Seidel and Kimble, 2011; Webster et al., 2022), so we hypothesized that pre-dauer nutrition may cause differences in the dauer reproductive system that impact post-dauer recovery of fertility. We discovered that *C. elegans* enter dauer with different numbers of germline cells depending on pre-dauer feeding. More severely starved worms have fewer cells and a smaller brood size upon recovery than those that fed more before dauer. Surprisingly, the DAF-2/IIS receptor and the DAF-7/TGF-β ligand were dispensable for this nutritional response. The onset of starvation rapidly diminished presentation of the stemness cue LAG-2 by the germline stem cell niche and induced a Notch-independent quiescent state in the germ line. These DTC and germline changes were reversed within hours of recovery from dauer, including exponential recovery of LAG-2 on the DTC that appears to be regulated post-translationally. Altogether, this work reports that differences in recovery of fertility after dauer and dauer gonad variation are mediated by pre-dauer nutrient access, with severely starved worms having small gonads and broods. Pre-dauer feeding affects niche and stem cell maintenance via dynamic regulation of the LAG-2 niche signal at the onset of and recovery from starvation.

## RESULTS

### Dauers that form in the presence of food have more germ cells and recover with larger broods than those that form after abject starvation

Different dauer induction regimes lead to differential reproductive recovery (Ow et al., 2018). We compared dauer gonads produced by various regimes. Two of these induce dauer under conditions of abject starvation: dauer entry after food depletion by mixed populations on parafilmed plates (Karp, 2018) and starvation in liquid culture of a synchronized population (Hibshman et al., 2021). Two other methods produce dauers in the presence of food using crowding (high density plating, (Ow and Hall, 2015)), or isolation of the first-formed dauers from crowded growth plates (this study, see Methods). We used an otherwise wild-type marker control strain expressing a germ cell nuclear marker (*mex-5p::H2B::mCherry::nos-2 3’UTR*) and an endogenously tagged integrin alpha subunit *ina-1(qy23[ina-1::mNG])* that aids visualization of the somatic gonad.

Worms that starved severely before dauer had about half to a third as many germ cells (average of 10.9 and 10.8 germ cells for starved plate and liquid culture, respectively) than those that had access to food (average of 22.2 and 37.8 germ cells for high density and first-formed, respectively, Fig. 1A,B), and we recapitulated the finding that dauers induced by high-density plating recover to have larger broods than those that starved before dauer (Fig. 1C). Brood size of first-formed dauers was also higher than starved dauers (Fig. 1C and Table S1A). Different dauer induction protocols changed overall gonad size but did not alter somatic cell numbers in the dauer gonad, only their proximity to one another (Fig. S1A-H), demonstrating that dauer gonads can vary in germ cell number and somatic cell size. The gonad displayed allometric scaling with body width and was relatively smaller in starved dauers compared to first-formed dauers (Fig. S1E), further demonstrating different growth dynamics in the soma and germ line.

**Figure 1:**
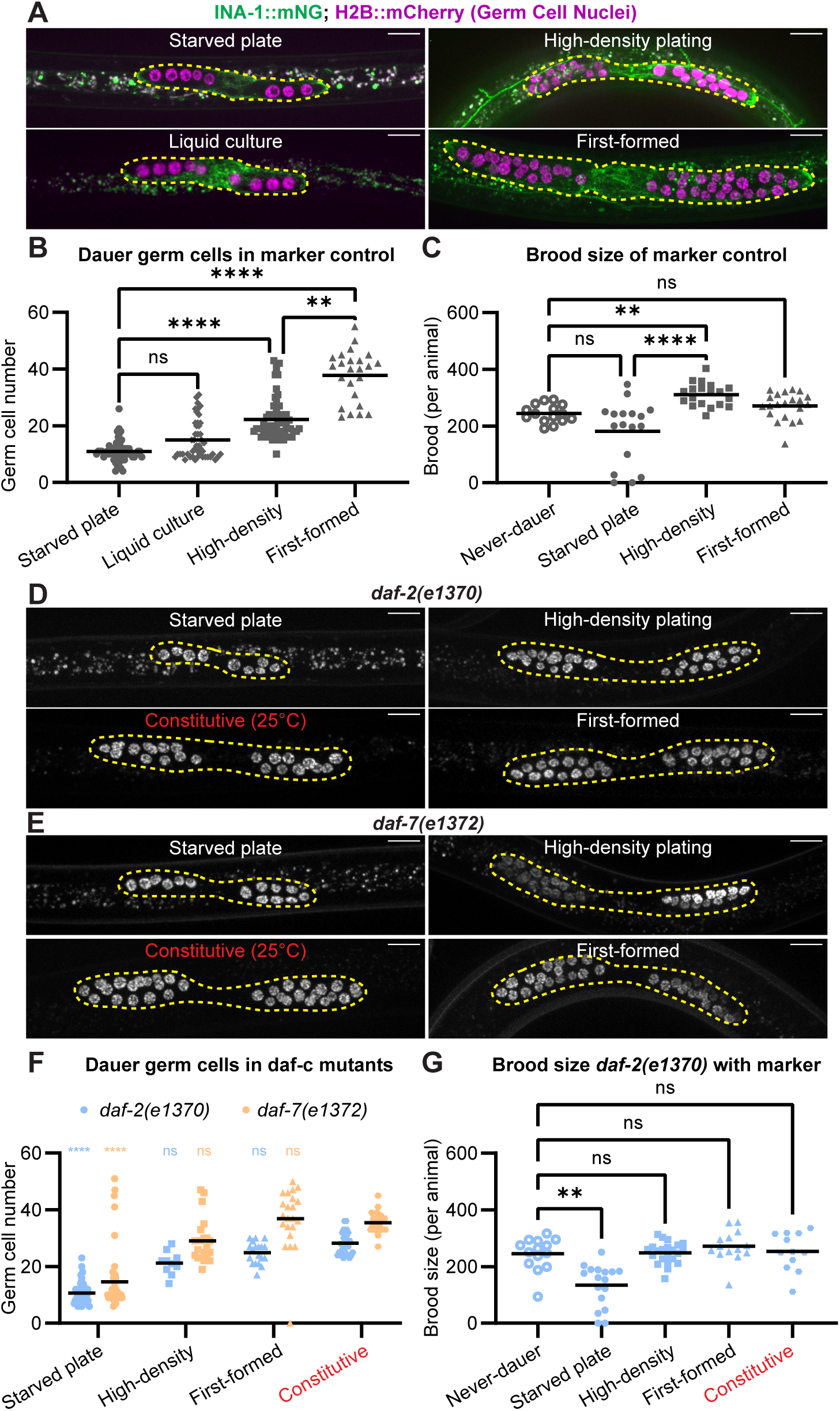
Dauer larvae that form after severe starvation have smaller gonads and recover to have smaller broods than dauer larvae that form after feeding. (A) Representative images of dauers induced with given methods. Gonads outlined. (B) Germ cell numbers in dauers of marker control strain induced by different methods. Starved plate (n=59), liquid culture (n=46), high density (n=61), and first-formed (n=24). Kruskal-Wallis test statistic H=111.0, p<0.0001. (C) Brood size of marker control strain. Never-dauer control (n=15), recovered dauers from starved plate (n=17), high-density plating (n=19), and first-formed (n=21). Kruskal-Wallis test statistic H=27.19, p<0.0001. (D-E) Representative fluorescence images of germ cells in dauers induced with given methods for (D) *daf-2(e1370)* and (E) *daf-7(e1372)* expressing germ cell nuclear marker. Gonads outlined. (F) Germ cell numbers in *daf-2(e1370)* and *daf-7(e1372)* dauers induced with four methods. Starved plate (n*_daf-2_* =52, n*_daf-7_* =47), high density (n*_daf-2_* =37, n*_daf-7_* =21), first-formed ( n*_daf-2_* =19, n*_daf-7_* =22), and constitutive (n*_daf-2_* =33, n*_daf-7_* =27). Kruskal-Wallis test statistic H=175.3 p<0.0001. (G) Brood size of *daf-2(e1370);naSi2(mex-5p::H2B::mCherry::nos-2 3’UTR)*. Never-dauer control (n=14), recovered dauers from starved plate (n=17), high density plating (n=15), first-formed (n=14), and constitutive late (n=12) recovered animals. Kruskal-Wallis test statistic=30.51, p<0.0001. All scale bars 10 μm. Dunn’s correction for multiple comparisons was used post-hoc to determine statistical significance of pairwise differences for relevant comparisons, asterisks on graph indicate these at p<0.0001 ****; p<0.005 **. Full results of statistical analyses in Supplemental Table S1.

We then used the field-standard dauer-formation constitutive (daf-c) mutants carrying reduction-of-function alleles encoding *daf-2(e1370)* (the sole IIS-like signaling receptor) or *daf-7(e1372)* (a TGF-β ligand), which enter dauer constitutively at high temperature even while feeding (Karp, 2018; Kenyon et al., 1993; Kimura et al., 1997; Ren et al., 1996; Thomas et al., 1993; Vowels and Thomas, 1992)). These are not thought to be true “temperature sensitive” mutations in which a gene product is functional at low temperature and nonfunctional at higher temperature, rather it is thought these are reduction-of-function alleles at all temperatures that are sensitized for dauer formation at high temperatures (Dr. Collin Ewald, ETH Zürich, personal communication, Ewald et al., 2018; Gems et al., 1998; Greer et al., 2008; Kenyon et al., 1993; McGehee, 2019; Trent et al., 1983; Tullet et al., 2008). High temperatures also increase the likelihood of dauer formation in wild-type worms (Ailion and Thomas, 2000). Into these genetic backgrounds we crossed the germ cell nuclear marker shown in Fig. 1A.

At low temperatures, daf-c mutants can form dauers using the same protocols that induce dauer in wild-type worms (Fig. 1D-F), and when they do, they display the same patterns as wild-type worms: few germ cells when severely starved before dauer, more germ cells when able to feed more before dauer (Fig. 1D-F). When *daf-2(e1370)* and *daf-7(e1372)* mutants are reared at high temperatures and constitutively form dauer despite feeding *ad libitum*, they have many germ cells (Fig. 1D-F). Thus daf-c worms, like wild-type worms, have different germ cell numbers in dauer depending on the dauer induction method.

Finally, we measured brood sizes of the *daf-2(e1370)* strain after recovery from dauer induced by different methods. Recovered, starvation-induced *daf-2(e1370)* dauers had lower brood size than recovered crowding-induced dauers, first-formed dauers, and constitutive dauers of that genotype, which all had the same brood size as the never-dauer control *daf-2(e1370)* (Fig. 1G). Our experiments implicate pre-dauer feeding as a potential determinant of germ cell number in dauer and brood size after recovery from dauer, and wild type alleles of neither *daf-2* nor *daf-7* are required for the response to pre-dauer feeding.

### The germ line grows exponentially with pre-dauer feeding in mutants with defective IIS and TGF-β signaling

To further test the hypothesis that pre-dauer feeding determines dauer germ cell number, we performed a series of food removal experiments on *daf-2(e1370)* and *daf-7(e1372)* daf-c mutants expressing the germ cell nuclear marker. Using daf-c mutants at 25°C ensured that we could vary food exposure while keeping every animal in the L2d dauer entry program. Worms were reared on plates with bacterial food for the specified number of hours at 25°C (Fig. 2A), removed from food at 25°C, and maintained until a population of majority dauers was noted (see Methods).

**Figure 2:**
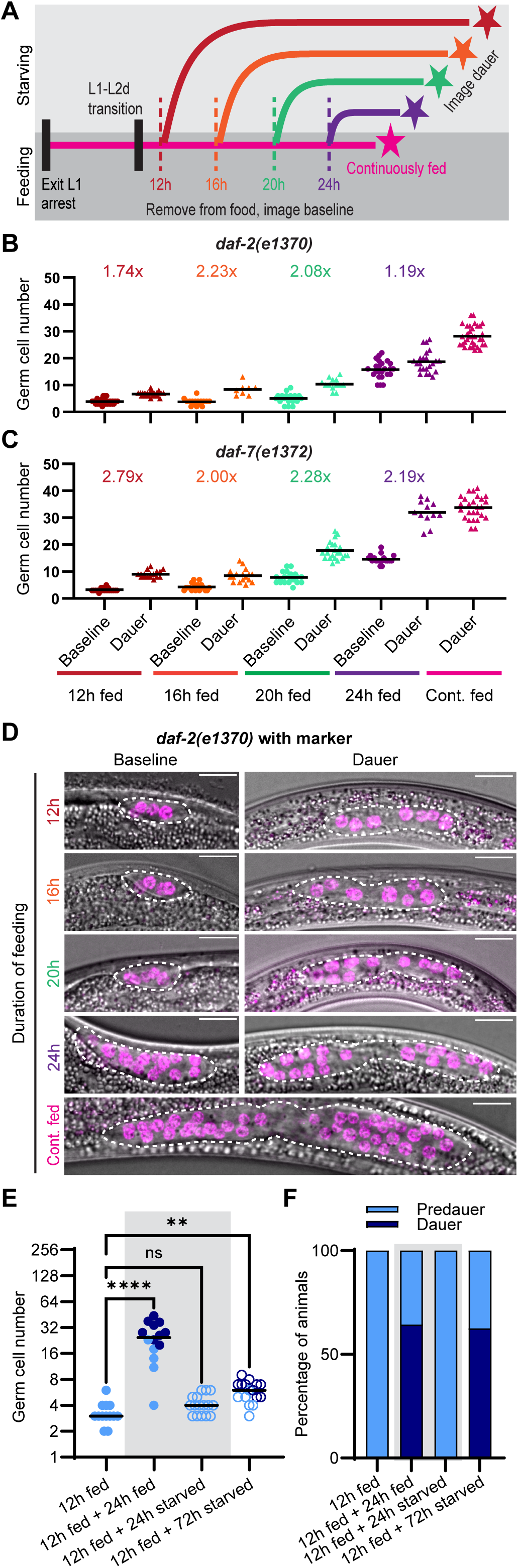
Withdrawal of food in the pre-dauer period arrests germ cell proliferation after a single round of division. (A) Schematic illustrating food removal experiments. Worms moved off food (dashed lines, imaged “Baseline” (2B-D)) and reared until dauer entry (stars, imaged “Dauer” (2B-D). All experiments in Fig. 2 were conducted at 25°C. Both strains express germ cell nuclear marker. (B) Results of experiment detailed in (2A), showing number of germ cell nuclei in *daf-2(e1370)*. Sample sizes: 12 h fed n_baseline_=31, n_dauer_=22; 16 h fed n_baseline_ =16, n_dauer_=8; 20 h fed n_baseline_=18, n_dauer_=13; 24 h n_baseline_=23, n_dauer_=24; continuously fed dauer n=33. (C) Results of experiment detailed in (2A), showing number of germ cell nuclei in *daf-7(e1372)*. Sample sizes: 12 h fed n_baseline_=20, n_dauer_=17; 16 h fed n_baseline_=20, n_dauer_ n=17; 20 h fed n_baseline_=20, n_dauer_=22; 24 h fed n_baseline_=18, n_dauer_=12; continuously fed dauer n=26. (D) Representative images of *daf-2(e1370)* measured for (2B) at time of food removal (baseline, left) and in dauer (right). Gonad outlined in white. *daf-7(e1372)* shown in Fig. S2. All scale bars 10 μm. (E) Experiment with *daf-2(e1370)* as in (2A) with additional +24 h endpoint. Darker dots indicate animals in dauer diapause at the time of counting (see 2E). n_12 h fed_=13; n_12 h fed, 24 h starved_=16; n_12 h fed, 24 h fed_ =14, refers to animals that were washed at 12 h and returned to food. n_12 h fed, 72 h starved_=16. Kruskal-Wallis test statistic=39.12; p<0.0001 with significance from Dunn’s correction for multiple comparisons indicated as p<0.0001 ****; p<0.005 **. (F) Percentage of worms that had reached the dauer stage under each condition in 2D. (E-F) Grey shaded box marks the samples quantified in parallel after 24 hours under experimental conditions, both on and off food.

We removed worms from food during the L2d stage at 12, 16, 20, 24 hours, and allowed others to feed continuously until dauer (48+ hours (Karp, 2018)). For each duration of feeding, we assayed germ cell number directly upon removal from food (Fig. 2 “Baseline”) and again within a day after dauer entry (Fig. 2 “Dauer”, cuticular alae visible, Fig. S2A,A’). These comparisons allowed us to determine the relationship between time spent feeding and germ cell number at both stages, and how much germ cell numbers changed between the cessation of feeding and dauer entry (Figs 2A-D, S2A). Baseline germ cell numbers increased with time spent feeding, and germ cell numbers at terminal gonad size in dauer increased exponentially for both *daf-2(e1370)* and *daf-7(e1372)* (Figs 2B,C, S2B,C). Thus we conclude it is not the different methods of inducing dauer that lead to differences in dauer gonad size, but the fact that worms feed more or less before entering dauer via these different methods.

While actively feeding (comparing among “Baseline” measures in Fig. 2B,C), the gonads of both *daf-2(e1370)* and *daf-7(e1372)* worms grow very little during the interval between 12 and 20 hours (Fig. 2B,C), however feeding during this stage sets up significant differences in terminal gonad size in dauer (for *daf-2(e1370)* mean_dauer fed 12 h_=6.68, mean_dauer fed 20 h_=10.38, Mann-Whitney U=16.50, p<0.0001; for *daf-7(e1372)* mean_dauer fed 12 h_=7.16, mean_dauer fed 20 h_=17.04, Welch’s t-test t=11.45, p<0.0001).

Comparing the rate of germline growth for *daf-2(e1370)* animals revealed an even more dramatic response to starvation than that of germ cell number alone (Fig. 2E). Animals gained an average of one germ cell per hour while feeding (12 h fed+24 h fed), while animals that starved (12 h fed+24 h starved) added on average only one germ cell total in that 24 hour period (Fig. 2E, gray shaded box) and entered dauer more slowly (Fig. 2F, gray shaded box). Our findings reveal pre-dauer germline responsiveness to nutrition is surprisingly robust to mutation in IIS and TGF-β signaling genes.

Average germ cell numbers approximately double in both genetic backgrounds between the food removal baseline and dauer entry at each timepoint (Fig. 2B,C), with the exception of 24 h of L2d feeding for *daf-2(e1370)* animals, which are known to slow germ cell division in late L2d in preparation for dauer entry (Narbonne and Roy, 2006). This slowing may also explain why terminal growth plateaus between 24 h dauers and continuously fed dauers. Germ cell doubling means cells undergo one additional round of division before dauer. Germ cells will complete an average of one additional cell division after DTC laser ablation in the L2-L3 stages of development (Kimble and White, 1981), and genetic ablation of the stemness receptor GLP-1/Notch allows adult germ cells to complete their current mitotic cell cycle to make a terminal division before differentiating (Fox and Schedl, 2015). Based on these established cell cycle constraints on stem-like germ cell proliferation dynamics, our finding that germ cells complete one round of division after food removal suggests that the withdrawal of food effectuates the rapid termination of pro-proliferative signaling to the L2d germ line. The pro-proliferative signal to the germ line is the DTC-expressed LAG-2 ligand, so we next asked whether this signal was down-regulated upon food removal.

### The DTC stemness cue LAG-2 requires *daf-2* and *daf-7* in fed L2s, but not for dramatic downregulation upon food removal in L2d

The pro-proliferative, anti-differentiation signal to the germ line is the DTC-expressed LAG-2 ligand activating GLP-1/Notch signaling in the distal germ cells (Henderson et al., 1994). We wanted to test whether shifting worms off food in the L2d sensitive window in which feeding changes germline growth (shown in Fig. 2) also alters the LAG-2 signal in the DTC. We used an endogenously tagged LAG-2::mNeonGreen and an integrated *lag-2* transcriptional reporter that drives a DTC membrane marker (*qIs154(lag-2p::myrTdTomato*) to quantify DTC LAG-2 abundance and *lag-2* transcriptional activity.

Otherwise wild-type dauer worms have notably less expression of both LAG-2::mNG and *lag-2* transcriptional reporter than fed L2 controls, though the magnitude of transcriptional reporter reduction in dauer (25-30%) cannot alone explain the reduction in LAG-2::mNG protein signal (∼30 fold) (Fig. 3A-D). Comparing otherwise wild-type worms to *daf-2(e1370)* and *daf-7(e1372)* reveals that both IIS and TGF-β are required for normal expression levels of of *lag-2* promoter activity and LAG-2::mNG protein in the DTC in fed L2 animals (Fig. 3E-I); this concords with what is known about the positive transcriptional regulation of the *lag-2* promoter by *daf-7* later in continuous, non-dauer development (Pekar et al., 2017).

**Figure 3:**
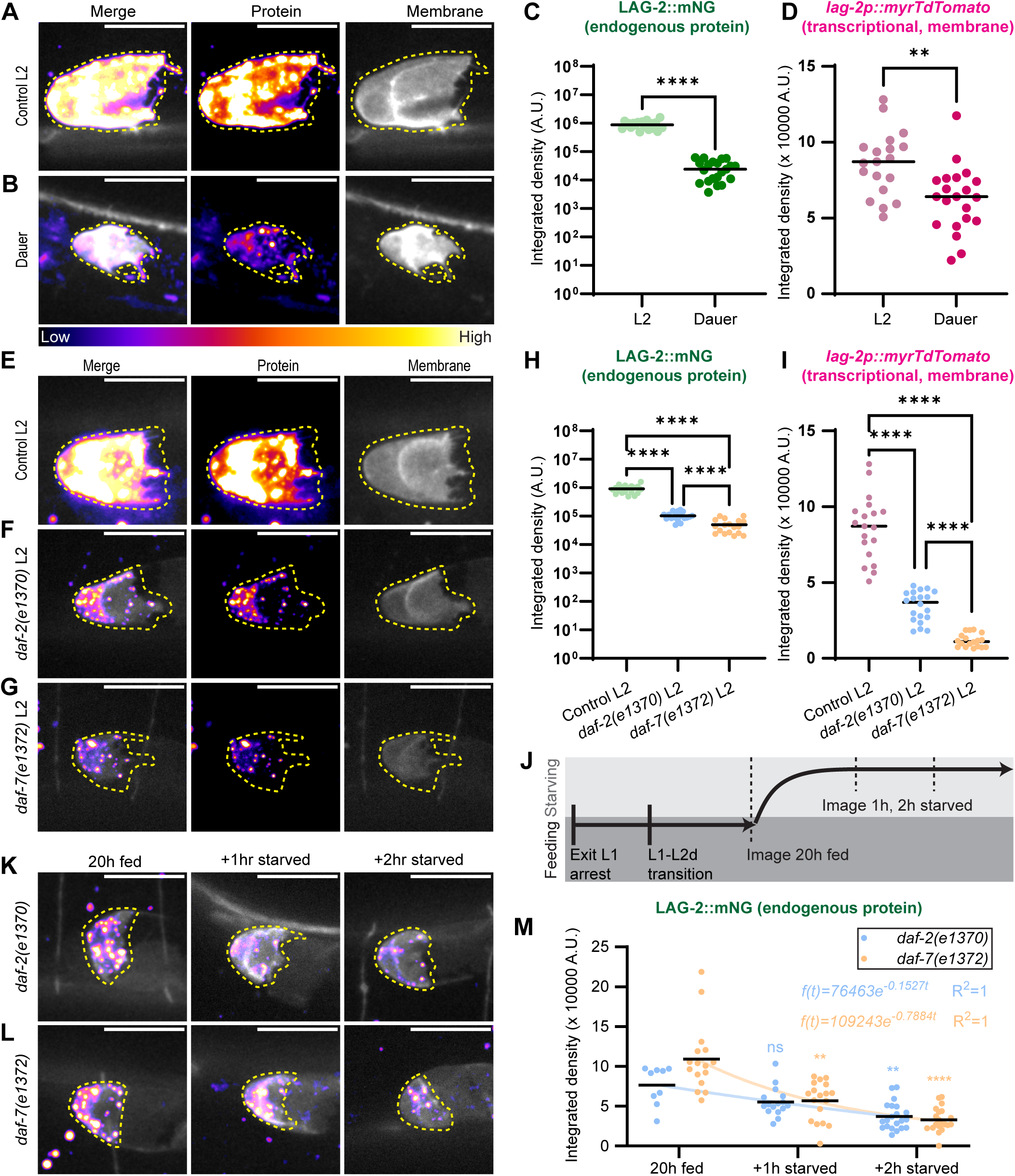
LAG-2 expression is higher in fed animals than in dauer and drops rapidly upon food removal in L2d, with *daf-2* and *daf-7* required for the former but not the latter. (A-B) Sum projections through eleven 0.3 µm z-slices of the superficial surface of a DTC expressing endogenously tagged LAG-2 protein (fire lookup table, *lag-2(cp193[lag-2:: mNeonGreen])*) and a membrane-localized transgenic reporter (grayscale, *qIs154(lag-2p:: myr::tdTomato)*) in fed L2 controls (n=19) and starved plate dauers (n=21). Each fluorescence channel is shown with identical scaling between treatments. (C-D) Integrated density measured for DTCs described in (A-B). (C) LAG-2::mNG, note log_10_ scale Mann-Whitney test statistic U=0, p<0.0001, and (D) *qIs154(lag-2p:: myr::tdTomato)*. Unpaired t-test with Welch’s correction t=3.491, p=0.0012. (E-G) Representative images of the transgenes in (A) in fed L2 worms that are (E) otherwise wild-type (n=19), (F) *daf-2(e1370)* mutants (n=20), (G) *daf-7(e1372)* mutants (n=18). (H-I) Integrated density measured for DTCs with sample sizes given in (E-G). (H) LAG-2::mNG, note log_10_ scale. Welch’s ANOVA test statistic W=89.99 (2.000, 18.76) p<0.0001 with Dunnett’s T3 multiple comparisons, and (I) *qIs154(lag-2p:: myr::tdTomato)*. Otherwise wild-type control, same L2 dataset as in (B-C). Welch’s ANOVA test statistic W=139.9 (2.000, 29.43) p<0.0001 with Dunnett’s T3 multiple comparisons. (J) Schematic of experiment performed for (K-M). Worms carrying the transgenes in the *daf-2(e1370)* and *daf-7(e1372)* mutant backgrounds were raised on food at 25°C for 20 h, imaged, moved off of food, and imaged one and two hours later. (K-L) Representative images of worms described in (J). Sum projection through eleven 0.3 µm z-slices of the superficial surface of the DTC at timepoints shown above for (K)*daf-2(e1370)* (n_20 h fed_=9, n_+1 h starved_=16, n_+2 h starved_=21) and (L) *daf-7(e1372)* mutants (n_20 h fe_=16, n_+1 h starved_=18, n_+2 h starved_=20) coexpressing endogenously tagged LAG-2 protein (fire lookup table) with DTC membrane-localized transcriptional reporter (grayscale). (M) Integrated density measured for LAG-2::mNG in DTCs with sample sizes given in K-L. Kruskal-Wallis test statistic H=53.78, p<0.0001 with Dunn’s correction for multiple comparisons; pairwise comparisons were made between 20 h fed and the two food-removal timepoints per genotype. Regression and R^2^ values displayed on graph. All scale bars 10 μm. DTCs outlined in yellow. Significance of pairwise comparisons after post-hoc corrections indicated as p<0.0001 ****; p<0.001 ***; p<0.01 **.

We could now compare expression of LAG-2::mNG for each daf-c genotype in a food removal assay in L2d (Fig. 3J). Endogenous LAG-2::mNG protein abundance is highly responsive to the onset of starvation in L2d in both genotypes (Fig. 3K-M). LAG-2::mNG drops from the L2d 20 h fed baseline within one hour of food removal in both *daf-2(e1370)* (-27.7%) and *daf-7(e1372)* (-48.1%) mutants, and continues to fall in an exponential decay (Fig. 3M). The abundance of most proteins is governed by first-order kinetics, leading to exponential decay in the absence of new protein synthesis ((McShane et al., 2016) and references therein). Thus our results suggest that LAG-2 deposition on the membrane in the DTC arrests nearly instantaneously upon food removal and existing protein is actively depleted thereafter. The starvation-induced arrest of germ cell proliferation that we observed in Fig. 2 is accompanied by a rapid starvation-induced downregulation of of LAG-2::mNG in the germline stem cell niche, and both processes are robust to deficits in *daf-2* and *daf-7* signaling, because we observe them in both reduction-of-function mutants.

This result distinguishes the process observed here from the *daf-7-*dependent downregulation of *lag-2* transcriptional reporters observed in L4 and young adult animals reared in unfavorable conditions (Dalfó et al., 2012; Pekar et al., 2017). Further comparisons of *lag-2* promoter activity and protein abundance in *daf-7(e1372)* mutants reinforced our conclusions that the role of *daf-7* in regulating *lag-2* in pre-dauer and dauer stages (Fig. S3) is distinct from the simple positive relationship reported during continuous development by Pekar et al. (2017). Across stages and conditions, *daf-7*/TGF-β signaling regulates *lag-2* in the DTC in a complex manner, but wildtype *daf-7* is not required for rapid downregulation of LAG-2 in the DTC after the acute onset of starvation (Fig. 3L,M).

### The dauer germ line is maintained in a Notch-independent, quiescent state, and exits that state within hours of dauer recovery

We next asked how the germ line tolerates diminished LAG-2 niche signal in dauer. Under continuous feeding, niche signaling via LAG-2 to the GLP-1/Notch receptor in the germ cells is required for their maintenance in a mitotic, undifferentiated state at all developmental stages (Austin and Kimble, 1989, 1987; Hansen and Schedl, 2006; Henderson et al., 1994; Kimble and Crittenden, 2007; Yochem and Greenwald, 1989). Both DTC ablation (Kimble and White, 1981) and genetic ablation of Notch signaling with a temperature-sensitive allele of *glp-1* (Cinquin et al., 2010; Fox and Schedl, 2015; Kodoyianni et al., 1992) are sufficient to trigger meiosis in all previously-mitotic distal germ cells within hours. However, transient adult starvation induces a G2-arrested, Notch-independent quiescent state in which genetic ablation of Notch signaling does not cause germ cells to differentiate; instead, germ cells remain mitotic upon subsequent recovery of Notch signaling at the permissive temperature (Seidel and Kimble, 2015).

Once in the dauer stage, worms are in a *de facto* starved state, and dauer germ cells are G2-arrested (Narbonne and Roy, 2006). We therefore hypothesized that the germ line of dauers, like that of transiently starved adults, is Notch-independent for the maintenance of an undifferentiated, stem-like population of germ cells.

We investigated dauer germ cell Notch-dependence using worms carrying a temperature sensitive allele of *glp-1*/Notch*, glp-1(bn18).* At the permissive temperature (16°C), *glp-1(ts)* mutants have adequate Notch signaling to remain fertile, but upon shifting to the restrictive temperature (25°C), they lose active Notch signaling and thus lose the germline stem cell population to differentiation (Kodoyianni et al., 1992). In adults, signs of meiotic entry appear in distal germ cells within six hours of temperature upshift (Fox and Schedl, 2015), so the germ cell fate response to lost Notch-signaling is rapid.

We first shifted continuously-fed control populations of *glp-1(bn18)* mutants to the restrictive temperature for 24 hours (green dashed path in Fig. 4A, and see Methods), then recovered single individuals of various larval stages to plates at the permissive temperature. Nearly all animals recovered as sterile adults (1/28 had a few offspring, Fig. 4B), demonstrating the expected Notch dependence of the fed larval germ line. If the germ line of dauer worms is likewise Notch-dependent, sterility should also occur when *glp-1(bn18)* dauers are shifted to the restrictive temperature.

**Figure 4:**
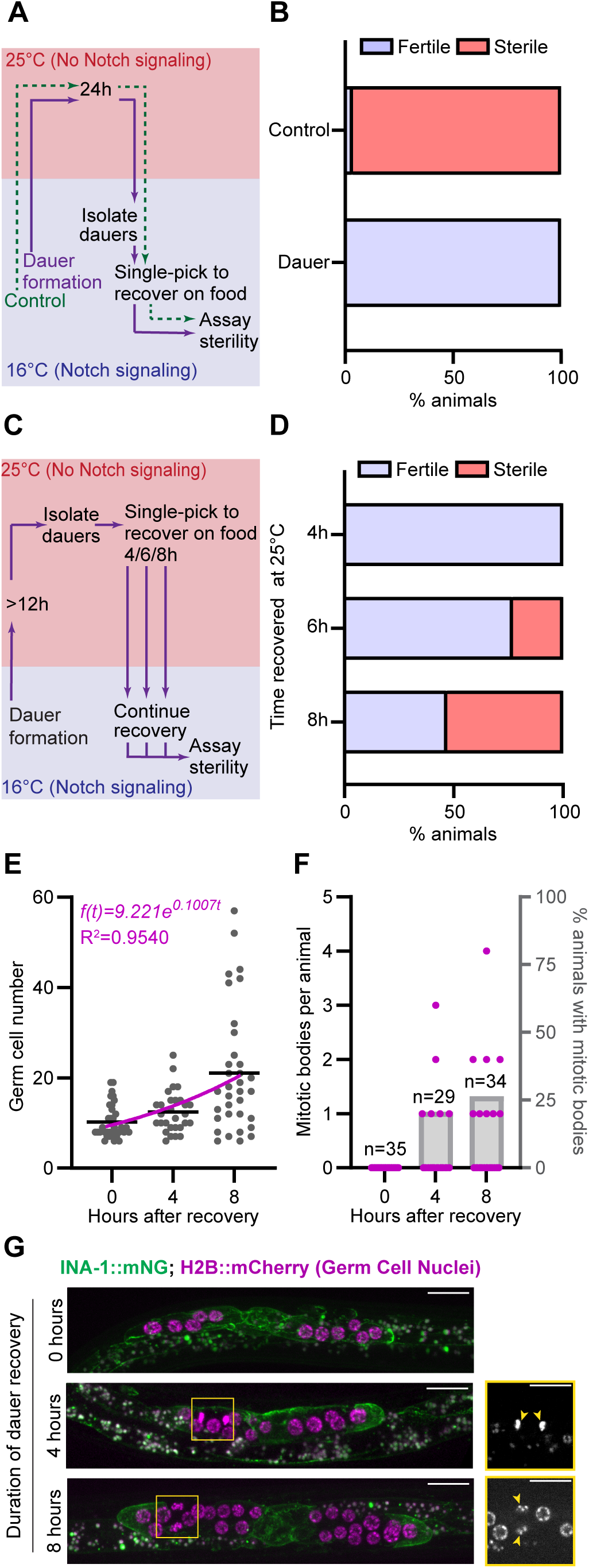
Dauer germ cells are arrested in a Notch-independent state and rapidly regain Notch dependence and cell cycling upon dauer exit. (A) Schematic illustrating temperature shift experiment testing germline *glp-1*/Notch dependence in dauer. (B) Worm fertility after (A). Control (never-dauer population temp shift in parallel with dauer) n=28, Recovered dauers n=29. (C) Schematic of dauer recovery *glp-1(bn18)* temperature-shift experiment testing the return of Notch dependence to the germ line. (D) Percentage of animals recovering as fertile vs. sterile results after recovery from dauer at 25°C for the first 4 h (n=15), 6 h (n=15) and 8 h (n=15) of dauer recovery. (E) Number of germ cells after specified hours of dauer recovery in the marker control strain n_0 h_=35, n_4 h_=29, n_8 h_=34. (F) Number of mitotic figures observed in each sample in E. (G) Representative images of gonads at 0 h, 4 h, and 8 h of recovery. Note 8 h sample has large and small nuclei typical of cells just before and after division.

This is not what we find. When we caused *glp-1(bn18)* mutant worms to form dauers at the permissive temperature and then shifted them to the restrictive temperature for 24 hours to abolish Notch signaling (purple solid path in Fig. 4A, and see Methods), isolated single dauer individuals to plates with food at the permissive temperature, and allowed the worms to recover, all of the worms recovered as fertile adults (n=29/29, Fig. 4B). Thus, the dauer germ line is in a Notch-independent, G2-arrested state and can maintain germline stem cells even in the absence of niche signaling. Because germ cells cannot directly transition from mitotic G2 to prophase I of the meiotic cell cycle without completing mitotic M phase (Fox and Schedl, 2015), cell-cycle quiescence protects dauer germ cells, like those in transiently starved adults (Seidel and Kimble, 2015), from differentiating in the absence of Notch activity.

We hypothesized that the germ line must return to a Notch-dependent state during dauer recovery. We performed temperature downshift experiments during dauer recovery for *glp-1(bn18)* mutant worms (Fig. 4C, and see Methods). When *glp-1(bn18)* mutant dauers were raised to 25°C, isolated, kept at that restrictive temperature for the first four hours of recovery from dauer, and then shifted to the permissive temperature to complete development, all animals recovered as fertile adults (Fig. 4D, n=15/15), meaning that germ cells remain Notch independent during early dauer recovery. After six hours of recovery from dauer at 25°C, a third of worms became sterile in adulthood (Fig. 4D, n=5/15). After eight hours of recovery from dauer at 25°C, more than half of worms recovered as sterile adults (Fig. 4D, n=8/15). The return of Notch dependence occurs at the earliest in the 4-6 hour window after recovery from dauer and is prevalent by eight hours of recovery.

Finally, we find that germ cell proliferation recommences in this same window, with a first doubling observed by eight hours, and modeled at ∼7 hours by our nonlinear regression (Fig. 4E). This proliferation rate accords with the conservative estimate of doubling in continuously fed L3 germ lines and those recovering from adult reproductive diapause (average of ∼9 hours, (Roy et al., 2016)). Mitotic figures reappear in the germ line during this time period (Fig. 4F,G). The germline mitotic index we observe (2.26% at four hours and 1.75% at eight hours) is intermediate between the mitotic indices reported for wild-type fed larvae and adults (Roy et al., 2016). The exit of G2-arrested quiescence and resumption of germ cell cycling temporally matches the return of Notch dependence in the germ line. We next asked when during recovery the Notch pathway ligand LAG-2 returns to high levels in the DTC.

### Dauer recovery initiates a transcription-independent burst of LAG-2 ligand presentation by the germline stem cell niche

We predicted that LAG-2 protein abundance would rapidly increase to meet the post-dauer requirement for germline Notch signaling. Immediately after dauer isolation (“0 h” of recovery, Fig. 5A-C), otherwise wild-type dauers had low LAG-2::mNG protein and *lag-2* transcriptional reporter signal. During the first two hours of dauer recovery, LAG-2::mNG protein abundance increased ∼4-fold, and continued to increase exponentially, reaching a 38-fold increase by eight hours post dauer recovery (Figs 5A,C, S4A, S5).

**Figure 5:**
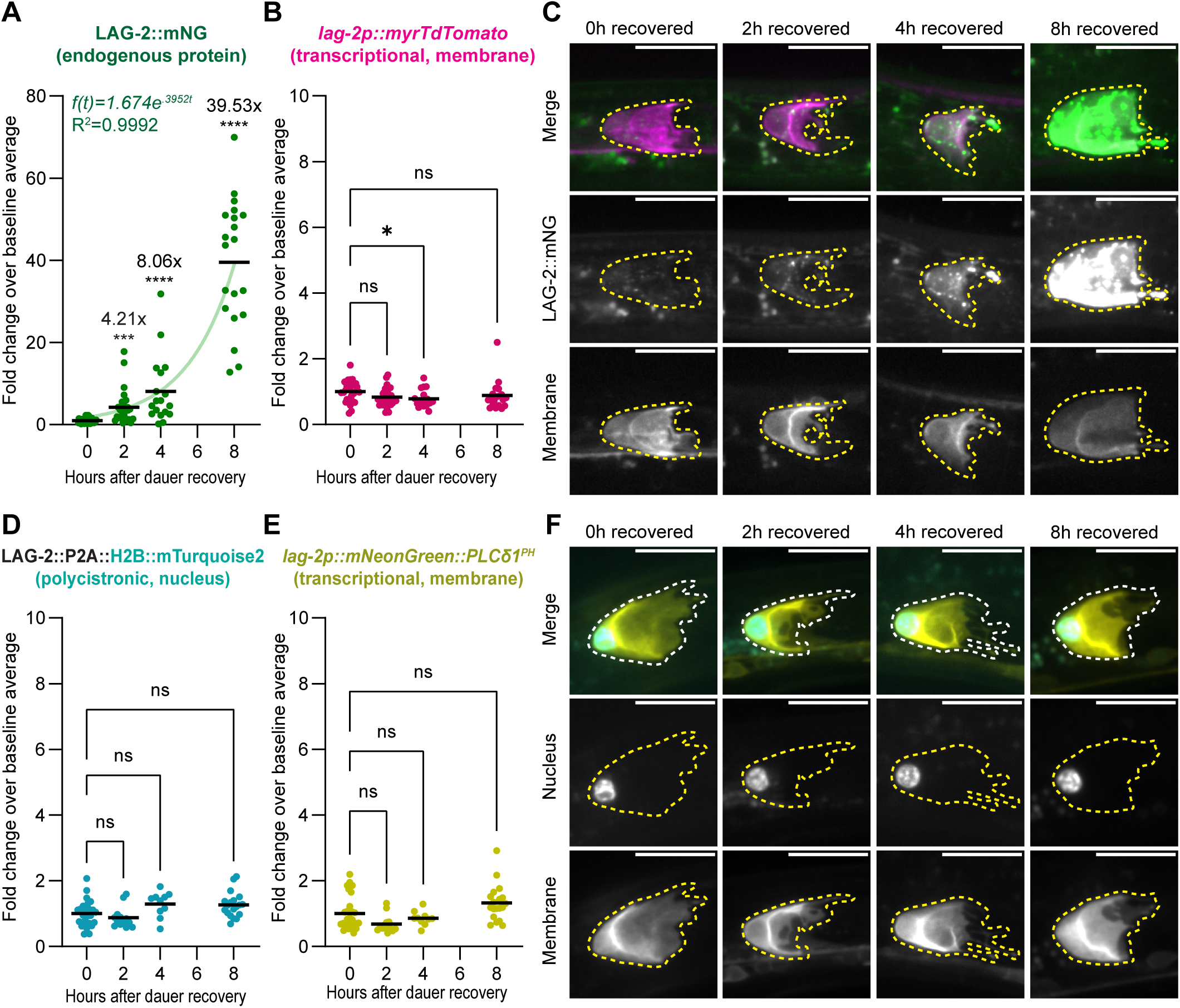
Recovery from dauer triggers a burst of LAG-2 presentation on the DTC niche without a coincident rise in *lag-2* transcription. (A-D) An otherwise wild-type strain coexpressing the endogenously tagged LAG-2 protein *lag-2(cp193[lag-2:: mNeonGreen])* (green in C) with a membrane-localized transgenic reporter *qIs154(lag-2p:: myr::tdTomato)* (magenta in C) recovering from dauer. (A) Fold change of LAG-2::mNG relative to mean at 0 h (n_0 h_ =30; n_2 h_ =29; n_4 h_=18; n_8 h_=20). Note Y-axis scale compared to other graphs. Kruskal-Wallis test statistic H=67.26, p<0.0001. Dunn’s correction for multiple comparisons show all pairwise comparisons are significantly different. (B) Fold change of the *lag-2p::myrTdTomato* transcriptional reporter for samples in A. Kruskal-Wallis test statistic H=8.905, p=0.0306. (C) Sum projection through eleven 0.3 µm z-slices of the superficial surface of a DTC for the time points shown above. Each fluorescence channel is shown with identical scaling between time points. (D-F) Otherwise wild-type strain co-expressing a *lag-2(bmd202* [*lag-2::P2A::H2B::mTurquoise2]*) polycistronic histone reporter knocked into the endogenous *lag-2* locus (blue in F) with a *lag-2p::mNeonGreen::PLCδ1^PH^* membrane-localized transcriptional reporter (yellow in F), same treatment as A-C. (D) Fold change of mTurquoise2 relative to mean at 0 h (n_0 h_ =29; n_2 h_=13; n_4 h_=10; n_8 h_=19). Kruskal-Wallis test statistic H=13.30m, p=0.0040, with Dunn’s correction for multiple comparisons showing that none of the recovery timepoints is significantly different from 0 h baseline. (E) Fold change of *lag-2p::mNeonGreen::PLCδ1^PH^* transcriptional reporter relative to mean at 0h (n_0 h_ =29; n_2 h_ =13; n_4 h_=10; n_8 h_=19). Kruskal-Wallis test statistic H=17.41, p=0.0006, with Dunn’s correction for multiple comparisons showing that none of the recovery timepoints is significantly different from 0 h baseline. (F) Sum projections through 0.3 µm Z-slices from superficial surface of DTC to bottom of nucleus. Non-normalized fluorescence measures and statistical analyses shown in Fig. S4. DTCs outlined. All scale bars 10 μm.

We did not observe a corresponding increase in signal from the *lag-2p::myrTdTomato* transcriptional reporter (Figs 5B,C, S4B), suggesting that the rapid restoration of LAG-2 protein on the DTC may not be regulated transcriptionally.

To further test this hypothesis, we examined another pair of *lag-2* reporters. To rule out potential artifacts caused by the multi-copy transgene or slow maturation of myrTdTomato fluorescence signal, we used a single-copy transcriptional reporter encoding a faster-folding fluorescent protein (*cpIs122(lag-2p::mNeonGreen::PLCδ1^PH^*). To rule out potential artifacts caused by the selection of the 3kb upstream promoter region for these transgenes and their lack of endogenous transcript regulation, we used a histone H2B::mTurquoise2 knocked into the endogenous *lag-2* locus with a P2A peptide cleavage site (*lag-2(bmd202*[*lag-2*::P2A::H2B::mTurquoise2^lox511I^2xHA])). This element will generate a polycistronic mRNA (Medwig-Kinney et al., 2022), each translation of which will produce one LAG-2 protein and one H2B::mTurquoise2 that separate at translation due to ribosome skipping at the P2A site (Donnelly et al., 2001; reviewed in de Lima and Lanza, 2021; Donnelly et al., 2001). The H2B::mTurquoise2 is therefore subject to endogenous *lag-2* transcript-based regulation (e.g. by transcriptional regulation, 3’ UTR-mediated translational repression by microRNAs or RNA binding proteins, transcript decay, etc.). RNAi against *lag-2* was used to validate this feature of the reagent (Fig. S6).

Up to eight hours after recovery from dauer, we observe no significant change in the levels of expression vs. baseline of either the faster-folding single copy transcriptional membrane reporter or polycistronic nuclear reporter (Figs 5D-F, S4C,D). Given that the germ line of 53% of animals is Notch dependent by eight hours of dauer recovery (Fig. 4D), transcription-based upregulation of *lag-2* would not appear to be sufficient to increase Notch ligand on the timeline necessary for dauer recovery.

Indeed, the only way to see an increase of endogenously tagged LAG-2::mNG but not an increase in the H2B::mTurquoise2 from the polycistronic knock-in is for the two proteins to be turned over at different rates. We ruled out that H2B::mTurquoise2 is particularly long-lived (see Methods and Fig. S7). This leaves only LAG-2 protein regulation as a source for their different recovery dynamics. We therefore conclude that the *lag-2* locus is transcriptionally active in the DTC during dauer–despite the germ line being maintained in a Notch-independent state–and LAG-2 is under protein-specific negative regulation that is rapidly alleviated upon dauer exit.

Taken together we see an exponential rebound of LAG-2 protein on the DTC begin within two hours of recovery from dauer, preceding a return of Notch dependence and proliferation in the germ line between 4-8 hours of recovery. Because dauer exit is asynchronous among individuals (Cassada and Russell, 1975), the precise timing of these events with respect to dauer exit and to one another may be even more tightly coordinated. Other early features of dauer exit–loss of the buccal plug/SDS resistance and resumption of pharyngeal pumping–also occur within a two-hour window of removal from dauer conditions (Cassada and Russell, 1975), making the return of LAG-2 on the DTC surface as rapid a response to the end of dauer as any other measured event. Our findings establish upregulation of germline stem cell niche signaling as a central, early feature of dauer recovery.

## DISCUSSION

The germ line is the only lineage in *C. elegans* with an indeterminate division pattern and variable cell number, while the somatic lineage is invariant (Kimble and Hirsh, 1979; Sulston and Horvitz, 1977; Sulston et al., 1983). This unique flexibility allows the germ line to be the only cell population that can respond to starvation by limiting the number of cells it produces, while the soma continues to the “safe harbor” state of dauer diapause. Starvation induces a precipitate IIS- and TGF-β-independent halt in pro-proliferative niche signaling and germ cell proliferation, which is swiftly reversible upon dauer recovery. That speed is particularly important as germ cells quickly exit the Notch independence of dauer and require active Notch signaling to maintain stemness as they resume cycling during recovery from dauer.

Rapid changes in protein levels and discordance of protein and transcriptional dynamics both suggest post-translational regulation (Liu et al., 2016). Exponential decay of LAG-2::mNG signal after food removal in L2d in daf-c worms suggests active degradation of LAG-2 without further production. The exponential recovery of LAG-2::mNG upon dauer exit suggests the cessation of that degradation process. We propose a model by which steady but fairly low levels of *lag-2* transcription and translation in dauer are coupled with LAG-2 protein degradation triggered by starvation. Upon dauer recovery, protein degradation ceases, leading to the rapid accumulation of LAG-2 protein in the DTC.

Such a regulatory paradigm famously governs the p53 family of tumor suppressor proteins, which are transcribed, translated, ubiquitinated, and degraded under normal growth conditions when p53 is not active. DNA damage or other stresses repress its ubiquitin ligase, allowing the rapid accumulation, nuclear translocation, and activity of p53 (Abuetabh et al., 2022). Recently, cyclin D has been discovered to be regulated by a similar process (Chaikovsky et al., 2021; Maiani et al., 2021; Simoneschi et al., 2021). Other proteins that mediate rapid response to changing environmental conditions are also regulated by steady state production and degradation in the absence of induction, like the hypoxia inducible transcription factor subunit HIF-α (reviewed in Flick and Kaiser, 2012 and in Weidemann and Johnson, 2008) and the Nrf2 transcriptional regulator of heme oxygenase HO-1 (Stewart et al., 2003; reviewed in Flick and Kaiser, 2012).

Notch signaling is regulated at the protein level in multiple ways (Fortini, 2009; Suarez Rodriguez et al., 2023; Tian et al., 2004; Wu et al., 2016), including by E3-ubiquitin-ligase (Fortini, 2009; Yamamoto et al., 2010). In *Drosophila* neuroblast stem cells, Notch signaling regulates a nutritionally-responsive entry into and exit from quiescence that is driven by amino acid availability, though this mechanism is insulin dependent (Sipe and Siegrist, 2017; Sood et al., 2022). Future work will investigate regulation of LAG-2 in the DTC by the ubiquitin-proteasome pathway, to test if it is regulated in a similar way to p53 upon starvation and during dauer exit. Such a regulatory mechanism “takes the foot off the brake” to rapidly induce a regulator under conditions in which its activity is required, priming a cell to respond to a high-stakes change in state. Protection of the germ line during starvation and recovery is a high-stakes regulatory event in the life of a worm.

Upon food removal, pre-dauer germ cells complete a single terminal division before dauer entry. The most severely starved dauers have as few germ cells (∼5-8) as worms with severe loss-of-function alleles of *glp-1*/Notch receptor gene (Austin and Kimble, 1987), and even fewer germ cells than worms have after early laser ablation of the DTC (Kimble and White, 1981), suggesting the cessation of pro-proliferative cues upon starvation is near-immediate. Thus we interpret the observation that germ cell number during recovery from dauer differs depending on how dauer was induced (Ow et al., 2021, 2018) to reflect differences in initial dauer germ cell number induced by these same treatments.

The question remains whether a reduction in DTC LAG-2 signal in response to starvation is the proximal cause of germline quiescence, or whether germline quiescence is a necessary precondition for the germ line to tolerate a reduction in DTC LAG-2 as a byproduct of starvation. Previous work reported that the dauer germ line is quiescent for germ cell division even with constitutively active Notch signaling (Narbonne and Roy, 2006), suggesting that downregulation of LAG-2 in the DTC is not necessary to prevent germ cell division in dauer itself, though the pre-dauer period we focus on here was not tested. Dauer germ cells are quiescent for proliferation and also for differentiation; meiotic germ cells in *glp-1(ts)* mutant dauers will not progress through spermatogenesis (Narbonne and Roy, 2006). This quiescence depends on AMPK signaling (*lbk-1* and *aak-2*) independent of the DAF-2 IIS receptor (Narbonne and Roy, 2006). The somatic gonad requires DAF-18/PTEN–independent of the DAF-2 IIS receptor or DAF-7/TGF-β–to enter a quiescent state and to mediate germline quiescence in dauer itself, but does not affect quiescence in the pre-dauer period that is the focus of our study (Tenen and Greenwald, 2019).

Expression of both endogenously tagged LAG-2 protein and *lag-2* promoter-driven transgenes (Fig. 5 and (Narbonne and Roy, 2006)) remain detectable in dauer, so the downregulation of *lag-2* during starvation is relative, not absolute. Downregulation of *lag-2* transcriptional reporters in response to starvation or pheromone occurs in later larval stages (Pekar et al., 2017), and the degree of that observed transcriptional difference (∼50%) is on the order of what we observe with transcriptional reporters in this study in dauers compared to fed L2s (∼30%). In contrast, LAG-2::mNG protein is reduced by *∼30 fold* in otherwise wild-type dauers compared to fed L2s and then increases by a similar magnitude within hours of recovery from dauer, a window of time in which the signal from transcriptional reporters does not rise at all. Notch signaling via *lag-2* is important in dauer and in germline regulation in many contexts (reviewed in Gordon, 2020). Expression of *lag-2* in the nervous system is required both during dauer and for dauer recovery (Ouellet et al., 2008), and *lag-2* reporter activation in the vulval precursor cells differs between continuous L2 vs. L2d and L3 vs. dauer (Karp and Greenwald, 2013). LAG-2 protein downregulation in the DTC has been observed during worm aging (Aprison et al., 2024 preprint; Singh et al., 2024), and *lag-2* transcription responds to male pheromone signals (Aprison et al., 2024 preprint). All levels of *lag-2* regulation–from transcriptional to post-translational–sculpt its dynamics in a range of cells to govern a number of processes, notably those regulating environmental response in the reproductive system.

While the precise molecular mechanisms linking nutrition to LAG-2 protein abundance in the DTC remain to be found, we establish that they proceed even with the field-standard *daf-7(e1372)* reduction of TGF-β signal and the *daf-2(e1370)* reduction of-function allele of the sole *C. elegans* IIS receptor. The reproductive consequences of pre-dauer food intake are also the same in *daf-2(e1370)* animals as they are in wild-type worms (that is, pre-dauer starvation leads to reduced brood size). In addition to seeking the molecular mechanisms of starvation-induced LAG-2 depletion in the DTC, future work will also aim to identify the systemic cues that trigger that depletion.

Early life stresses are known to affect later life outcomes in a range of organisms and contexts with potential mechanisms ranging from epigenetics to inflammation to hormone signaling to lasting variation in central nervous system development and function (Taylor, 2010). In *C. elegans,* primordial germ cells are kept in a quiescent state by the somatic gonad precursors (McIntyre and Nance, 2023). Arrested vs. fed L1 worms have differential transcriptional responses in the soma and germ line (Webster et al., 2022), and display chromatin compaction in the germ line (Belew et al., 2021; Morao and Ercan, 2021). The L1 arrest state is triggered by complete starvation after hatching (Baugh and Hu, 2020; Baugh, 2013; Johnson et al., 1984) and several checkpoints triggered by later starvation can arrest somatic development (Baugh and Hu, 2020; Schindler et al., 2014).

Future work will focus on how the germ line transitions from a Notch-independent state back to Notch-fueled germline proliferation. One possible driver is nucleotide levels, which have been shown to limit cell division but not cell growth in unicellular organisms (Diehl et al., 2022) and to affect germline growth in non-dauer *C. elegans* (Chi et al., 2016). Considering nucleotide levels and the limits imposed by starvation upon cellular energy prompts the observation that continuous transcription, translation, and LAG-2 protein degradation during dauer comes with an energy cost. A dauer worm would only pay this price if the alternative–slower restoration of LAG-2 protein to the germline stem cell niche surface–carried an even greater cost of reduced fertility upon return of the germ line to Notch dependence.

## MATERIALS AND METHODS

Sections of this text are adapted from prior Gordon lab publications (Li et al., 2022; Singh et al., 2024), as they describe our standard laboratory practices and equipment.

### Strains

We used Wormbase (Sternberg et al., 2024) while conducting this study. Some strains were provided by the CGC, which is funded by NIH Office of Research Infrastructure Programs (P40 OD010440) and are to be requested directly from CGC. The following *C. elegans* strains were obtained from the CGC: N2 (Brenner, 1974), CB1370 *daf-2(e1370)* III (Kenyon et al., 1993; Kimura et al., 1997), CB1372 *daf-7(e1372)* III (Pierce et al., 2001; Ren et al., 1996), DG2389 *glp-1(bn18)* III (Kodoyianni et al., 1992).

The following *C. elegans* strains were generously shared by community members or generated previously in our lab (with sources given for each strain and allele): NK2517 (Gordon et al., 2019) *qIs154(lag-2p:: myr::tdTomato) (Byrd et al., 2014); lag-2(cp193[lag-2:: mNeonGreen^3xFlag])* V)

KLG034 (Gordon et al., 2019) *cpIs122(lag-2p::mNeonGreen:: PLCδ1^PH^)*II (Linden et al., 2017)*; lag-2(bmd202* [*lag-2*::P2A::H2B::mTurquoise2^lox511I^2xHA]) V (Medwig-Kinney et al., 2022),

GS9692 *arTi435(rps-27p::2xnls::gfp(flexon)::unc-54 3′ UTR* I); *arTi237(ckb-3p::Cre(opti)::tbb-2 3′ UTR* X) (Shaffer and Greenwald, 2022).

The following *C. elegans* strains were generated for this paper by crossing existing alleles and markers (with sources given for alleles not previously attributed above): KLG047 *ina-1(qy23[ina-1::mNeonGreen]) (Jayadev et al., 2019)*; *naSi2*(*mex-5p::H2B::mCherry::nos-2 3’UTR)* II (Linden et al., 2017). We call this the “marker control strain” in the text and refer to the *nasi2* transgene as “germ cell histone marker”.

KLG050 *daf-2(e1370)* III*; naSi2(mex-5p::H2B::mCherry::nos-2 3’UTR)* II, KLG051 *daf-2(e1370)* III*; qIs154(lag-2p:: myr::tdTomato)* V*; lag-2(cp193[lag-2:: mNeonGreen^3xFlag])* V.

KLG055 *daf-7(e1372)* III*;nasi2*(*mex-5p::H2B::mCherry::nos-2 3’UTR)* II, KLG056 *daf-2(e1372)* III*; qIs154(lag-2p:: myr::tdTomato)* V*; lag-2(cp193[lag-2:: mNeonGreen^3xFlag])* V.

### Strain maintenance and synchronization

Worm strains were maintained on nematode growth media (NGM) at 16°C unless otherwise specified.

### Dauer formation by starvation on parafilmed plates

To more precisely time the formation of facultative dauers, a large, mixed population of well-fed worms was transferred from an uncrowded growth plate to NGM plates seeded with ∼70 μL of an overnight culture of *E. coli* OP50 bacterial food. Plates were sealed with parafilm and placed at 25°C for the non-temperature-sensitive strains and 16°C for strains containing *daf-2(e1370), daf-7(e1372),* or *glp-1(bn18)* temperature-sensitive alleles. Plates were monitored daily for evidence of starvation (bacterial lawn depleted and worms dispersed). At 25°C this took ∼2-4 days and at 16°C this took ∼5-7 days. Plates were analyzed after 1+ weeks of starvation.

### Recovery of first-formed dauers

As above, evaluated <4 days after food exhaustion was noted, before a robust population of dauers are visible on the plate. Dauers were isolated with a treatment of 1% SDS for 20 minutes (see Dauer isolation, below).

### Dauer formation by liquid culture

For dauer formation by liquid culture, the protocol from Hibshman et al., 2021 was used. Briefly, animals were egg prepped (Stiernagle, 2006) from 4-5 well-populated NGM plates seeded with OP50 and embryos were allowed to hatch out in S-complete medium overnight such that a synchronized population of L1s was generated. Animals were concentrated to 5 worms/μL in S-complete and added to glass test-tubes or 25 mL Erlenmeyer flasks, to which *E. coli* HB101 bacteria was added from a concentrated stock such that the final concentration of bacteria in S-complete was 1 mg/mL. Cultures were placed on a shaker (180 rpm). Deviating from the original protocol, animals were incubated at 25°C for 4-5 days and then treated with SDS to match the dauer isolation protocol used in our other methods.

### Dauer formation by high density plating

For high-density plating conditions, the protocol described in Ow & Hall 2015 (Ow and Hall, 2015) and summarized by (Karp, 2018) was used. Briefly, mixed populations of worms were chunked to four 100 mm plates seeded with 1 mL *E. coli* OP50 or 6-8 60 mm plates seeded with the same titre of food. Plates were grown until densely populated but not depleted of bacterial food. Worms were washed from plates in M9 buffer and allowed to gravity-settle, after which the supernatant (containing younger worms) was removed from the pellet of adults, which was retained for transfer to a single 35 mm plate seeded with *E. coli* OP50. We seeded the 35 mm plates with more bacterial food (100-200 uL of 20x concentrated *E. coli* OP50) than the original protocol (50 uL of an overnight culture) to ensure that the animals had abundant food available prior to dauer formation. The culture plates were then blanketed with a cooked, pasteurized egg white mixture as per the original protocol. To these plates, the pelleted adults were added. For daf-c strains at 16°C, plates were incubated for longer than the 72 h specified by the original protocol, as worm development is slower at that temperature. To recover dauers, plates were SDS-treated when there were dauers visible in a sample of the egg white mixture.

### Dauer isolation protocols

For plates with mixed populations, dauers were isolated with 1% SDS (Karp, 2018). After SDS treatment, the population of recovered animals was placed onto an unseeded NGM plate and the excess liquid was allowed to dry, after which animals were picked to a slide for imaging, were kept on the unseeded plate for the specified number of hours to evaluate short-term recovery, or picked to an NGM plate seeded with OP50 for recovery and brood measurement. For synchronous daf-c populations in dauer, worms were picked from all-dauer populations and dauer morphology was ascertained for each specimen by microscopy.

### Daf-c mutant constitutive dauer formation

Constitutive dauer formation of daf-c mutants (*daf-2(e1370)* and *daf-7(e1372)*) was achieved by collecting embryos via a standard egg prep (Stiernagle, 2006) with embryos hatching into L1s in M9 to create a synchronized population. This population was concentrated and added to NGM plates seeded with *E. coli* OP50. The mutant daf-c strains used in this study were the reduction of function alleles of the insulin-like receptor, class II allele *daf-2(e1370),* and the TGF-ꞵ ligand, *daf-7(e1372)* that are field-standard genetic backgrounds for studying dauer (Karp, 2018).

### Brood size assays

All brood size assays were conducted at 16°C to match the necessary conditions of the daf-c strains. After experimental treatment (see below), worms were singly picked to NGM plates seeded with ∼70 μL *E. coli* OP50. Egg-laying adults were subsequently passaged to a new plate once daily until egg laying stopped. Plates with progeny were kept at 16°C until the oldest worm on the plate had reached L4, after which they were counted. Brood totals include broods of animals that died during recovery and animals that were fully sterile, does not include animals that failed to recover from dauer to adulthood.

For never-dauer control brood assays, fed L4 animals of each strain were singled from mixed populations on Day 0. For brood after recovery from constitutive dauer formation, healthy plates of adults were egg prepped; eggs were rolled in M9 overnight at room temperature to obtain a population of arrested L1 larvae. Animals were dropped onto NGM plates seeded with OP50. Plates were sealed with parafilm and put at the restrictive temperature (25°C) to induce dauer formation. Worms were then isolated by SDS treatment (see next section) and dauers were singly recovered to NGM plates seeded with OP50 at 16°C to recover to reproductive adulthood.

Brood size was not measured for *daf-7(e1372)* mutant strains because these worms have a high rate of bagging, even at the permissive temperature (Karp, 2018; Shaw et al., 2007; Trent et al., 1983).

### Food removal experiments

A synchronized population of *daf-2(e1370); naSi2(mex-5p::H2B::mCherry::nos-2 3′UTR II)* L1s or *daf-7(e1372); naSi2(mex-5p::H2B::mCherry::nos-2 3′UTR II)* was obtained via an egg prep of at least eight 60 mm NGM plates seeded with *E. coli OP50*.

Synchronized L1s were added to NGM plates seeded with OP50 and were put at 25°C to induce constitutive dauer formation. After the specified number of hours (Fig. 2A), animals were rinsed off the seeded plate with M9, recovered to a 15 mL conical tube, and were subsequently washed in M9 3-5 times and then rolled in an excess of M9 at room temperature for 20 minutes before being washed an additional time in M9 to clear bacteria from the skin and gut. To circumvent worms sticking to the side of the conical tube, tubes were briefly vortexed between spins as needed.

Washed animals were transferred to peptone-free plates (recipe from (Eustice et al., 2022)) and returned to 25°C to continue to develop. Imaging was done immediately after animals were washed off the food plates (baseline), and parallel populations were kept at 25°C until the plate visibly had >50% dauers (the rest being pre-dauers; these strains do not progress past dauer at 25°C). Animals that appeared to be dauers under a dissecting scope were picked to image for the dauer endpoint of the experiment; only animals with dauer morphology (recognized by radial constriction and the presence of alae) were measured for germ cell number. While removal of food in the L1 stage prior to the L1-L2d transition caused L1 developmental arrest, as expected, we found that removal of food after the L2d molt did not inhibit *daf-2(e1370)* or *daf-7(e1372)* animals from progressing through L2d or entering dauer.

At this temperature, these strains experience a ∼100% developmental arrest in the dauer stage and will not molt beyond dauer unless returned to lower temperatures (Gems et al., 1998; Karp, 2018; Kenyon et al., 1993; Ren et al., 1996). We used this endpoint rather than a set clock-time because the duration of the pre-dauer period of daf-c mutants at 25°C is much longer than the duration of the L1 and L2 stages of fed animals. The *daf-2(e1370)* mutant takes approximately 80 hours to enter dauer, and *daf-7(e1372)* takes approximately 48 hours (Karp, 2018), compared to just over 24 hours for continuously fed animals to reach L3 (Byerly et al., 1976). The pre-dauer period is also extended in wild-type animals that feed on a lower titre of bacterial food (Cassada and Russell, 1975).

### *glp-1(ts)* temperature shift experiments

Temperature shift experiments used the temperature-sensitive allele of the Notch receptor *glp-1(bn18)*. Loss of Notch signaling causes failure of germline induction (if the temperature shift happens in the embryonic period) or irreversible loss of the stem-like cell fate and meiotic entry of all germ cells (Fox and Schedl, 2015; Kodoyianni et al., 1992). To test for *glp-1* dependence during dauer (Fig. 4A,B), two populations were used. As a control, an unstarved, uncrowded population was reared at 16°C, shifted to the restrictive temperature of 25°C for 24 hours, larval worms were singly picked to NGM plates seeded with *E. coli* OP50, and these plates were kept at 16°C until adulthood. The experimental group was made of worms that had been starved on plates at 16°C to form dauers, which were shifted to 25°C for 24 hours. After 24 hours, dauers were recovered by SDS isolation and singled to NGM plates seeded with *E. coli* OP50 and kept at 16°C until adulthood. Each plate was subsequently assayed for the presence of progeny.

To test for when *glp-1* dependence returns after dauer (Fig. 4C,D), *glp-1(bn18)* worms were put into dauer on starved plates at 16°C (n=14 first formed, and n=15 starved), shifted to 25°C for a minimum of 12 hours, SDS isolated to trigger dauer exit, and transferred to NGM plates with food and placed back at 25°C for an additional four, six, or eight hours. At that time, the developmentally oldest/largest worms on the plate (the worms that exited dauer first, since dauer exit is asynchronous) were picked as single animals to NGM plates seeded with *E. coli* OP50 food and kept at 16°C until adulthood. After four days of recovery at 16°C, each plate was assayed for live larval offspring, and those that had offspring were scored as fertile. Parental worms on plates that lacked larvae were imaged at 60x magnification and scored for the presence of embryos in the uterus (fertile) or an absence of gametes (sterile). Embryo retention appeared to be a phenotype of *glp-1(bn18*).

### Confocal imaging

All images were acquired at room temperature on a Leica DMI8 with an xLIGHT V3 confocal spinning disk head (89 North) with a 63× Plan-Apochromat (1.4 NA) objective and an ORCAFusion GenIII sCMOS camera (Hamamatsu Photonics) controlled by microManager. RFPs were excited with a 555 nm laser; GFP and mNG were excited with a 488 nm laser; mTurquoise was excited with a 445 nm laser. Z-stacks through the gonad were acquired with a z-step size of 0.3 or 0.5 µm as noted. Worms were mounted on agar pads in M9 buffer with 0.01-0.02 M sodium azide paralytic (VWR (Avantor) Catalog Number 26628-22-8). Samples used for fluorescence quantification were acquired with the same laser power, exposure time, and z-step size within the datasets to be compared, with histograms monitored to ensure proper exposure.Images with poor body placement, debris on the slide, or other image quality concerns were not analyzed.

### Image analysis software

Images were processed in FIJI89 (Version: 2.14.1/1.54f).

### Dauer recovery experiments

To measure fluorescent protein expression and germ cell proliferation during dauer recovery, worms were isolated as described above (Dauer isolation protocol), and subsequently split such that some worms were imaged immediately (“0 hours of recovery”, Figs 4,5), and other worms were recovered to *E. coli* OP50-seeded NGM plates for the specified length of time.

### Germ cell proliferation during dauer recovery

Our otherwise wild-type marker control strain coexpressing *ina-1(qy23[ina-1::mNeonGreen]); naSi2*(*mex-5p::H2B::mCherry::nos-2 3’UTR)* II was put into dauer by starvation on plates, and dauers were isolated as above (Dauer isolation protocol).

Worms were imaged at that 0 h starting point, and parallel populations were recovered to plates seeded with *E. coli* OP50 bacterial food. Worms were imaged at four and eight hours after plating on food. Germ cells were counted manually and scored for mitotic figures (metaphase and anaphase chromatin condensations visible in the H2B::mCherry signal, Fig. 4G. We plotted germ cell numbers over time and the percentage of worms for which we observed mitotic figures in Fig. 4F. Mitotic indices were calculated as number of mitotic cells/total cells for each specimen as in (Roy et al., 2016) and averages for each timepoint are reported in the text.

### Fluorescence intensity measurement of reporters of *lag-2*: Endogenously tagged LAG-2::mNG with coexpressed membrane TdTomato

Strain NK2517 *qIs154(lag-2p:: myr::tdTomato); lag-2(cp193[lag-2:: mNeonGreen^3xFlag]) V* was used to measure endogenously tagged LAG-2::mNG protein (Gordon et al., 2019) and a coexpressed multicopy integrated transcriptional reporter of the *lag-2* ∼3 kb upstream promoter with an *unc-54* 3’ UTR (Byrd et al., 2014). All worms, including controls, were reared at 25°C. TdTomato has a half-time to maturation of one hour at 37°C (Shaner et al., 2004), while mNeonGreen has a half-time to maturation of fewer than ten minutes at 37°C (Shaner et al., 2013) (both will fold slower at worm-rearing temperatures). TdTomato signal is visible in adjacent neurons (Fig. 3 and Fig. 5) and the autofluorescent gut granules of the worm are visible in the channel of the endogenously tagged LAG-2::mNG. These structures were avoided as much as possible during projection, tracing, and measurement of signal.

We chose an L2 control for this experiment because, while the dauer larva is an alternative third larval stage, the worm’s reproductive system undergoes morphogenetic changes and changes in cell number during the normal L3 stage. These developmental events are arrested in dauer (Tenen and Greenwald, 2019), making L3 a confounding control for the dauer larvae reproductive system.

Confocal z-stacks were acquired with a 0.3 µm step size. A sum intensity z-projection was created in FIJI consisting of eleven slices representing the superficial half of the cell (from the superficial cell surface to the cross-section of the nucleus, which is visible as a dark void in the center of the cell body). Usually one DTC from each worm was analyzed, as the gut often obscures one. The DTC was hand-traced on the membrane marker RFP channel; that ROI was subsequently used to measure both the RFP (membrane) and GFP (endogenously tagged protein) channels, and slid off the DTC to the adjacent body of the worm to measure the background for both channels. Background subtracted Integrated Density measurements were thus obtained. We then normalized all integrated density measurements to the mean integrated density at the 0 h timepoint for that fluorescent protein, so fold-change values and statistics are shown in Fig. 5A,B,D,E, with raw values shown in Fig. S4. Kruskal-Wallis test with follow up Dunn’s correction for multiple comparisons of the raw, non-normalized integrated densities show the same pattern of significance (that is, only LAG-2::mNG signal increases relative to baseline) as the fold change analysis (Fig. S4). Integrated density was used for this measure instead of raw integrated density so that size differences in the DTC did not influence our measurements, as we noted that dauer DTCs are smaller than non-dauer L2 DTCs (Fig. S1I-J). To display the ∼40x range of LAG-2::mNG intensity with the same image scaling, some pixels are saturated in display images (Fig.5C). Saturation was not observed in the raw images from which measurements were made (Fig. S5).

### Fluorescence intensity measurement of reporters of *lag-2*: Polycistronic histone reporter with coexpressed membrane mNeonGreen

We considered the ∼3 kb upstream *lag-2* promoter fragment used in the transcriptional reporter transgenes could insufficiently capture endogenous transcriptional regulation, or that post-transcriptional regulation of *lag-2* mRNA could uniquely affect LAG-2 production. The membrane-localized transcriptional reporters that we use from (Byrd et al., 2014; Linden et al., 2017) like those of (Blelloch et al., 1999; Henderson et al., 1994; Pekar et al., 2017) are driven by ∼3 kb upstream of the *lag-2* transcription start site (including the TGF-β-responsive element described by (Pekar et al., 2017). This ∼3 kb element is sufficient to drive expression in DTCs, however (Karp and Greenwald, 2013) identified regulatory elements of *lag-2* as far away as ∼6.4 kb upstream, raising the possibility that ∼3 kb *lag-2* reporters are missing relevant regulatory information that is important for DTC expression under certain, previously unexamined conditions.

We therefore examined a histone H2B::mTurquoise2 (half-time to maturation of 33.5 minutes, (Goedhart et al., 2012) that was knocked into the endogenous *lag-2* locus with a P2A peptide (Medwig-Kinney et al., 2022). This element will generate a polycistronic mRNA, the translation of which will produce one LAG-2 protein and one H2B::mTurquoise2, meaning that the histone reporter is under the same transcriptional and post-transcriptional, transcript-based regulation (e.g. by 3’ UTR-mediated repression by microRNAs or RNA binding proteins, transcript decay, etc.) as the endogenous *lag-2* gene. It is coexpressed with another DTC-expressed transcriptional reporter, a faster-folding, membrane-localized *lag-2p::mNeonGreen::PLCδ1^PH^* driven by the ∼3 kb upstream *lag-2* promoter with a *let-858* 3’ terminator sequence, integrated into the MoscI site on Ch. II (*cpIs122, (Linden et al., 2017)*). This strain was previously analyzed in (Singh et al., 2024) and (Li and Gordon, 2025).

Confocal z-stacks were acquired with a 0.3 µm step size. A sum intensity z-projection through a 6-7 µm depth of the DTC (to capture the entire nucleus in z, ∼20 slices) was generated in FIJI, the DTC was hand-traced, and membrane fluorescence was measured for the GFP channel in that ROI. A background measurement was made by sliding the DTC ROI onto the adjacent body of the worm. The nuclear histone signal was obtained using an ellipse tool tight around the nucleus in the same z-projection to measure on the CFP channel, and background for the nuclear signal was made by sliding that ROI onto the adjacent body of the worm. Background subtracted Integrated Density measurements were thus obtained. We then normalized all integrated density measurements to the mean integrated density at the 0 h timepoint for that fluorescent protein, so fold-change values and statistics are shown in Fig. 5A,B,D,E, with raw values shown in Fig. S4. Kruskal-Wallis test with follow up Dunn’s correction for multiple comparisons of the raw, non-normalized integrated densities show the same pattern of significance (that is, no pairwise difference relative to baseline for either fluorescent protein) as the fold change analysis (Fig. S4).

To test whether perdurance of H2B::mTurquoise2 could mask the addition of new protein upon transcriptional upregulation (Fig. 5D), we examined its dynamics in the Z1 and Z4 lineages in early larval stages (Fig. S7). Late L1 and early L2 worms (12 h after release from L1 arrest at 25°C), and later L2 (15 h) worms were imaged as above and sum projections were made through ∼3.3 microns (eleven slices) of the superficial gonad arm (and for one sample, for both arms). This captured the DTC (Z1.aa or Z4.pp) and its sister cell (Z1.ap or Z4.pa), or else caught the parent of these cells (Z1.a or Z4.p) right before it divides. We did not note any systematic difference between Z1 and Z4-lineage cells for expression of this marker despite notably different membrane fluorescence in these cells driven by the ∼3 kb upstream *lag-2* promoter (Singh et al., 2025). For each gonad arm, we used an elliptical ROI to measure H2B::mTurquoise2 signal in the nuclei of these cells, as well as a background region in the middle of the gonad where no nucleus expresses the marker. The nucleus of the sister cell was identified for measurement as a void in the GFP channel if no histone signal could easily be noted. Background-subtracted measurements were made and compared to quantify perdurance of the fluorescent histone signal.

### lag-2 RNAi

To verify that the H2B::mTurquoise2 is indeed coregulated at the transcript level with endogenous *lag-2*, we measured the effect of *lag-2* RNAi on H2B::mTurquoise2 signal (Fig. S6). Single-colony overnight cultures were made of *E. coli* HT115 containing either the empty vector L4440 or clone Y73C8B.4 from the Ahringer library (Kamath and Ahringer, 2003; Kamath et al., 2003), which is complementary to >900 nt of the *lag-2* mRNA extending from just upstream of the start codon through exon 1, intron 1, and some of exon 2. Cultures contained ampicillin (100 μg/mL, VWR (Avantor), Catalog no. 76204-346) and were grown at 37°C, expression was induced with 1 mM IPTG (Apex BioResearch Products, cat# 20-109) for 1 h at 37°C, and the culture was plated and allowed to grow at least overnight at room temperature on NGM plates prepared with IPTG and ampicillin. L2 larval stage worms were picked onto NGM plates without food and allowed to move around for at least 30 minutes to prevent bacterial carryover. Animals were then hand-picked either onto *lag-2* RNAi or empty L4440 vector control plates, cultured for 24 h at 16°C to the L4 stage, imaged, and background-subtracted fluorescence intensity was measured as above.

### Somatic gonad marker measurements

*Somatic gonad cell number (Fig. S1G):* The strain GS9692 expressing *arTi435(rps-27p::2xnls::gfp(flexon)::unc-54 3′ UTR)*; *arTi237(ckb-3p::Cre(opti)::tbb-2 3′ UTR)* (Shaffer and Greenwald, 2022) was specifically designed to activate enduring GFP expression in the cells of the somatic gonad. We manually counted gonad cells in this strain after inducing first-formed and starved plate dauers (Fig. S1F-H). We always see the expected two DTCs and 10 gonad blast cells in dauers of any age. Variation in expression levels/tissue depth can sometimes complicate detection of the cell at the center of the gonad primordium (more orange cells in Fig. S1G). Landmarks were selected based on relative position with DTCs at the gonad tips and the next-most-distal fluorescent cells being the SS cells used for measuring.

### Statistical analysis

All statistical analysis was done using GraphPad Prism version 10.4.2 for Windows, GraphPad Software, Boston, Massachusetts USA, www.graphpad.com. All tests and sample sizes are specified in the relevant figure legends.

Before any statistical tests were conducted on data, the data was first evaluated for normal distribution using the Shapiro-Wilk test for normality. Statistical tests were selected based on the appropriate test given the normality or non-normality of the data. If the data were normally distributed, Welch’s t-test (two groups) or Welch’s ANOVA (three or more groups) was used because it is more robust against Type I error rates than Student’s t-test or an ordinary one-way ANOVA (Delacre et al., 2019; Derrick et al., 2016). When data violated the assumption of normality, non-parametric Mann-Whitney (two groups) or Kruskal-Wallis (three or more groups) tests were used. Post-hoc analyses for pairwise comparisons were selected based on appropriateness for the test used.

For curve fitting in Figs 3M, 4E, 5A, S2B,C, nonlinear regression models were compared to null-hypothesis linear regression models to test which was a better fit. Curves were fit to the means of the data at each timepoint. For exponential decay of LAG-2 protein, the plateau of the model was not constrained.

## Author Contributions

Conceptualization: F.A.K. and K.L.G.; Formal analysis: F.A.K.; Investigation: F.A.K., C.P.M., Data collection assistance: B.K.; Writing – original draft preparation: F.A.K. and K.L.G.; Writing – review and editing: F.A.K., C.P.M., and K.L.G.; Visualization: F.A.K.; Supervision: K.L.G.; Project administration: K.L.G.; Funding acquisition: F.A.K. and K.L.G.

## Acknowledgements

We thank members of the Gordon Lab and F.A.K. dissertation committee members R.H.D. and D.R.M. for thoughtful comments on the manuscript.

## Data and resource availability

All relevant data and details of resources can be found within the article. All *C. elegans* strains are available upon request, except those sourced from the CGC (above).

## Supplemental Material

**Table S1.** Statistical analyses for Fig. 1.

**Figure S1.** Somatic gonad cell number is unaffected by method of dauer induction.

**Figure S2.** *daf-7* and *daf-2* are dispensable for nutritionally-sensitive exponential gonad growth before dauer.

**Figure S3.** Expression dynamics of *lag-2* transcriptional reporter and LAG-2::mNG protein vary across stages in *daf-2(e1370)* and *daf-7(e1372)* mutants.

**Figure S4.** Non-normalized expression of LAG-2::mNG and *lag-2* reporters during dauer recovery show no increase in transcriptional activity.

**Figure S5.** Image scaling for display of qualitative comparison creates image saturation that is not present in the original data.

**Figure S6.** Knockdown of *lag-2(bmd202[lag-2::P2A::H2B::mTurquoise2])* signal by *lag-2* RNAi verifies H2B::mTurquoise2 transcript-based coregulation with *lag-2*.

**Figure S7.** Signal decay of *lag-2(bmd202* [*lag-2::P2A::H2B::mTurquoise2])* reveals effective detection of continuous vs. previous expression.

## References

Abuetabh, Y., Wu, H.H., Chai, C., Al Yousef, H., Persad, S., Sergi, C.M., Leng, R., 2022. DNA damage response revisited: the p53 family and its regulators provide endless cancer therapy opportunities. Exp. Mol. Med. 54, 1658–1669. 10.1038/s12276-022-00863-4.

Ailion, M., Thomas, J.H., 2000. Dauer formation induced by high temperatures in Caenorhabditis elegans. Genetics 156, 1047–1067. 10.1093/genetics/156.3.1047.

Angelo, G., Van Gilst, M.R., 2009. Starvation protects germline stem cells and extends reproductive longevity in C. elegans. Science 326, 954–958. 10.1126/science.1178343.

Aprison, E.Z., Dzitoyeva, S., Ruvinsky, I., 2024. The roles of TGFβ and serotonin signaling in regulating proliferation of oocyte precursors and germline aging. bioRxiv.

Austin, J., Kimble, J., 1989. Transcript analysis of glp-1 and lin-12, homologous genes required for cell interactions during development of C. elegans. Cell 58, 565–571. 10.1016/0092-8674(89)90437-6.

Austin, J., Kimble, J., 1987. glp-1 is required in the germ line for regulation of the decision between mitosis and meiosis in C. elegans. Cell 51, 589–599. 10.1016/0092-8674(87)90128-0.

Barbieri, M., Bonafè, M., Franceschi, C., Paolisso, G., 2003. Insulin/IGF-I-signaling pathway: an evolutionarily conserved mechanism of longevity from yeast to humans. Am. J. Physiol. Endocrinol. Metab. 285, E1064–71. 10.1152/ajpendo.00296.2003.

Baugh, L.R., Hu, P.J., 2020. Starvation responses throughout the caenorhabditiselegans life cycle. Genetics 216, 837–878. 10.1534/genetics.120.303565.

Baugh, L.R., 2013. To grow or not to grow: nutritional control of development during Caenorhabditis elegans L1 arrest. Genetics 194, 539–555. 10.1534/genetics.113.150847.

Belew, M.D., Chien, E., Wong, M., Michael, W.M., 2021. A global chromatin compaction pathway that represses germline gene expression during starvation. J. Cell Biol. 220. 10.1083/jcb.202009197.

Blelloch, R., Anna-Arriola, S.S., Gao, D., Li, Y., Hodgkin, J., Kimble, J., 1999. The gon-1 gene is required for gonadal morphogenesis in Caenorhabditis elegans. Dev. Biol. 216, 382–393. 10.1006/dbio.1999.9491.

Brenner, S., 1974. THE GENETICS OF *caenorhabditis elegans*. Genetics 77, 71–94. 10.1093/genetics/77.1.71.

Byerly, L., Cassada, R.C., Russell, R.L., 1976. The life cycle of the nematode Caenorhabditis elegans. I. Wild-type growth and reproduction. Dev. Biol. 51, 23–33. 10.1016/0012-1606(76)90119-6.

Byrd, D.T., Knobel, K., Affeldt, K., Crittenden, S.L., Kimble, J., 2014. A DTC niche plexus surrounds the germline stem cell pool in Caenorhabditis elegans. PLoS ONE 9, e88372. 10.1371/journal.pone.0088372.

Cassada, R.C., Russell, R.L., 1975. The dauerlarva, a post-embryonic developmental variant of the nematode Caenorhabditis elegans. Dev. Biol. 46, 326–342. 10.1016/0012-1606(75)90109-8.

Chaikovsky, A.C., Li, C., Jeng, E.E., Loebell, S., Lee, M.C., Murray, C.W., Cheng, R., Demeter, J., Swaney, D.L., Chen, S.H., Newton, B.W., Johnson, J.R., Drainas, A.P., Shue, Y.T., Seoane, J.A., Srinivasan, P., He, A., Yoshida, A., Hipkins, S.Q., McCrea, E., Sage, J., 2021. The AMBRA1 E3 ligase adaptor regulates the stability of cyclin D. Nature 592, 794–798. 10.1038/s41586-021-03474-7.

Chi, C., Ronai, D., Than, M.T., Walker, C.J., Sewell, A.K., Han, M., 2016. Nucleotide levels regulate germline proliferation through modulating GLP-1/Notch signaling in C. elegans. Genes Dev. 30, 307–320. 10.1101/gad.275107.115.

Cinquin, O., Crittenden, S.L., Morgan, D.E., Kimble, J., 2010. Progression from a stem cell-like state to early differentiation in the C. elegans germ line. Proc Natl Acad Sci USA 107, 2048–2053. 10.1073/pnas.0912704107.

Claeys, I., Simonet, G., Poels, J., Van Loy, T., Vercammen, L., De Loof, A., Vanden Broeck, J., 2002. Insulin-related peptides and their conserved signal transduction pathway. Peptides 23, 807–816. 10.1016/s0196-9781(01)00666-0.

Corsi, A.K., Wightman, B., Chalfie, M., 2015. A Transparent Window into Biology: A Primer on Caenorhabditis elegans. Genetics 200, 387–407. 10.1534/genetics.115.176099.

Dalfó, D., Michaelson, D., Hubbard, E.J.A., 2012. Sensory regulation of the C. elegans germline through TGF-β-dependent signaling in the niche. Curr. Biol. 22, 712–719. 10.1016/j.cub.2012.02.064.

Das, D., Arur, S., 2017. Conserved insulin signaling in the regulation of oocyte growth, development, and maturation. Mol. Reprod. Dev. 84, 444–459. 10.1002/mrd.22806.

Delacre, M., Leys, C., Mora, Y.L., Lakens, D., 2019. Taking Parametric Assumptions Seriously: Arguments for the Use of Welch’s *F*-test instead of the Classical *F*-test in One-Way ANOVA. rips 32, 13. 10.5334/irsp.198.

Derrick, B., Toher, D., White, P., 2016. Why Welch’s test is Type I error robust. TQMP 12, 30–38. 10.20982/tqmp.12.1.p030.

de Lima, J.G.S., Lanza, D.C.F., 2021. 2A and 2A-like Sequences: Distribution in Different Virus Species and Applications in Biotechnology. Viruses 13. 10.3390/v13112160.

Diehl, F.F., Miettinen, T.P., Elbashir, R., Nabel, C.S., Darnell, A.M., Do, B.T., Manalis, S.R., Lewis, C.A., Vander Heiden, M.G., 2022. Nucleotide imbalance decouples cell growth from cell proliferation. Nat. Cell Biol. 24, 1252–1264. 10.1038/s41556-022-00965-1.

Donnelly, M.L.L., Luke, G., Mehrotra, A., Li, X., Hughes, L.E., Gani, D., Ryan, M.D., 2001. Analysis of the aphthovirus 2A/2B polyprotein “cleavage” mechanism indicates not a proteolytic reaction, but a novel translational effect: a putative ribosomal “skip”. J. Gen. Virol. 82, 1013–1025. 10.1099/0022-1317-82-5-1013.

Erkut, C., Kurzchalia, T.V., 2015. The C. elegans dauer larva as a paradigm to study metabolic suppression and desiccation tolerance. Planta 242, 389–396. 10.1007/s00425-015-2300-x.

Eustice, M., Konzman, D., Reece, J.M., Ghosh, S., Alston, J., Hansen, T., Golden, A., Bond, M.R., Abramowitz, L.K., Hanover, J.A., 2022. Nutrient sensing pathways regulating adult reproductive diapause in C. elegans. PLoS ONE 17, e0274076. 10.1371/journal.pone.0274076.

Ewald, C.Y., Castillo-Quan, J.I., Blackwell, T.K., 2018. Untangling Longevity, Dauer, and Healthspan in Caenorhabditis elegans Insulin/IGF-1-Signalling. Gerontology 64, 96–104. 10.1159/000480504.

Flick, K., Kaiser, P., 2012. Protein degradation and the stress response. Semin. Cell Dev. Biol. 23, 515–522. 10.1016/j.semcdb.2012.01.019.

Fortini, M.E., 2009. Notch signaling: the core pathway and its posttranslational regulation. Dev. Cell 16, 633–647. 10.1016/j.devcel.2009.03.010.

Fox, P.M., Schedl, T., 2015. Analysis of Germline Stem Cell Differentiation Following Loss of GLP-1 Notch Activity in Caenorhabditis elegans. Genetics 201, 167–184. 10.1534/genetics.115.178061.

Gems, D., Sutton, A.J., Sundermeyer, M.L., Albert, P.S., King, K.V., Edgley, M.L., Larsen, P.L., Riddle, D.L., 1998. Two pleiotropic classes of daf-2 mutation affect larval arrest, adult behavior, reproduction and longevity in Caenorhabditis elegans. Genetics 150, 129–155. 10.1093/genetics/150.1.129.

Gimond, C., Poullet, N., Vielle, A., Demoinet, E., Braendle, C., 2025. Developmental plasticity of hermaphrodite sperm production across environments in Caenorhabditis elegans. PLoS ONE 20, e0336162. 10.1371/journal.pone.0336162.

Goedhart, J., von Stetten, D., Noirclerc-Savoye, M., Lelimousin, M., Joosen, L., Hink, M.A., van Weeren, L., Gadella, T.W.J., Royant, A., 2012. Structure-guided evolution of cyan fluorescent proteins towards a quantum yield of 93%. Nat. Commun. 3, 751. 10.1038/ncomms1738.

Golden, J.W., Riddle, D.L., 1984. The Caenorhabditis elegans dauer larva: developmental effects of pheromone, food, and temperature. Dev. Biol. 102, 368–378. 10.1016/0012-1606(84)90201-X.

Gordon, K.L., Payne, S.G., Linden-High, L.M., Pani, A.M., Goldstein, B., Hubbard, E.J.A., Sherwood, D.R., 2019. Ectopic Germ Cells Can Induce Niche-like Enwrapment by Neighboring Body Wall Muscle. Curr. Biol. 29, 823–833.e5. 10.1016/j.cub.2019.01.056.

Gordon, K., 2020. Recent Advances in the Genetic, Anatomical, and Environmental Regulation of the C. elegans Germ Line Progenitor Zone. J. Dev. Biol. 8. 10.3390/jdb8030014.

Greer, E.R., Pérez, C.L., Van Gilst, M.R., Lee, B.H., Ashrafi, K., 2008. Neural and molecular dissection of a C. elegans sensory circuit that regulates fat and feeding. Cell Metab. 8, 118–131. 10.1016/j.cmet.2008.06.005.

Hall, S.E., Beverly, M., Russ, C., Nusbaum, C., Sengupta, P., 2010. A cellular memory of developmental history generates phenotypic diversity in C. elegans. Curr. Biol. 20, 149–155. 10.1016/j.cub.2009.11.035.

Hand, S.C., Denlinger, D.L., Podrabsky, J.E., Roy, R., 2016. Mechanisms of animal diapause: recent developments from nematodes, crustaceans, insects, and fish. Am. J. Physiol. Regul. Integr. Comp. Physiol. 310, R1193–211. 10.1152/ajpregu.00250.2015.

Hansen, D., Schedl, T., 2006. The Regulatory Network Controlling the Proliferation–Meiotic Entry Decision in the Caenorhabditis elegans Germ Line, in: Current Topics in Developmental Biology. Elsevier, pp. 185–215. 10.1016/S0070-2153(06)76006-9.

Harsimran Kaur Gill, Gaurav Goyal, Gurminder Chahil, 2017. Insect diapause: A review. JAST-A 7. 10.17265/2161-6256/2017.07.002.

Henderson, S.T., Gao, D., Lambie, E.J., Kimble, J., 1994. lag-2 may encode a signaling ligand for the GLP-1 and LIN-12 receptors of C. elegans. Development 120, 2913–2924. 10.1242/dev.120.10.2913.

Hibshman, J.D., Webster, A.K., Baugh, L.R., 2021. Liquid-culture protocols for synchronous starvation, growth, dauer formation, and dietary restriction of Caenorhabditis elegans. STAR Protocols 2, 100276. 10.1016/j.xpro.2020.100276.

Hirsh, D., Oppenheim, D., Klass, M., 1976. Development of the reproductive system of Caenorhabditis elegans. Dev. Biol. 49, 200–219. 10.1016/0012-1606(76)90267-0.

Jayadev, R., Chi, Q., Keeley, D.P., Hastie, E.L., Kelley, L.C., Sherwood, D.R., 2019. α-Integrins dictate distinct modes of type IV collagen recruitment to basement membranes. J. Cell Biol. 218, 3098–3116. 10.1083/jcb.201903124.

Johnson, T.E., Mitchell, D.H., Kline, S., Kemal, R., Foy, J., 1984. Arresting development arrests aging in the nematode Caenorhabditis elegans. Mech. Ageing Dev. 28, 23–40. 10.1016/0047-6374(84)90150-7.

Kamath, R.S., Ahringer, J., 2003. Genome-wide RNAi screening in Caenorhabditis elegans. Methods 30, 313–321. 10.1016/s1046-2023(03)00050-1.

Kamath, R.S., Fraser, A.G., Dong, Y., Poulin, G., Durbin, R., Gotta, M., Kanapin, A., Le Bot, N., Moreno, S., Sohrmann, M., Welchman, D.P., Zipperlen, P., Ahringer, J., 2003. Systematic functional analysis of the Caenorhabditis elegans genome using RNAi. Nature 421, 231–237. 10.1038/nature01278.

Karp, X., Greenwald, I., 2013. Control of cell-fate plasticity and maintenance of multipotency by DAF-16/FoxO in quiescent Caenorhabditis elegans. Proc Natl Acad Sci USA 110, 2181–2186. 10.1073/pnas.1222377110.

Karp, X., 2018. Working with dauer larvae. WormBook 2018, 1–19. 10.1895/wormbook.1.180.1.

Kenyon, C., Chang, J., Gensch, E., Rudner, A., Tabtiang, R., 1993. A C. elegans mutant that lives twice as long as wild type. Nature 366, 461–464. 10.1038/366461a0.

Kimble, J., Crittenden, S.L., 2007. Controls of germline stem cells, entry into meiosis, and the sperm/oocyte decision in Caenorhabditis elegans. Annu. Rev. Cell Dev. Biol. 23, 405–433. 10.1146/annurev.cellbio.23.090506.123326.

Kimble, J., Crittenden, S.L., 2005. Germline proliferation and its control. WormBook 1–14. 10.1895/wormbook.1.13.1.

Kimble, J., Hirsh, D., 1979. The postembryonic cell lineages of the hermaphrodite and male gonads in Caenorhabditis elegans. Dev. Biol. 70, 396–417. 10.1016/0012-1606(79)90035-6.

Kimble, J.E., White, J.G., 1981. On the control of germ cell development in Caenorhabditis elegans. Dev. Biol. 81, 208–219. 10.1016/0012-1606(81)90284-0.

Kimura, K.D., Tissenbaum, H.A., Liu, Y., Ruvkun, G., 1997. daf-2, an insulin receptor-like gene that regulates longevity and diapause in Caenorhabditis elegans. Science 277, 942–946. 10.1126/science.277.5328.942.

Kim, S., Paik, Y.-K., 2008. Developmental and reproductive consequences of prolonged non-aging dauer in Caenorhabditis elegans. Biochem. Biophys. Res. Commun. 368, 588–592. 10.1016/j.bbrc.2008.01.131.

Klass, M., Hirsh, D., 1976. Non-ageing developmental variant of Caenorhabditis elegans. Nature 260, 523–525. 10.1038/260523a0.

Kodoyianni, V., Maine, E.M., Kimble, J., 1992. Molecular basis of loss-of-function mutations in the glp-1 gene of Caenorhabditis elegans. Mol. Biol. Cell 3, 1199–1213. 10.1091/mbc.3.11.1199.

Linden, L.M., Gordon, K.L., Pani, A.M., Payne, S.G., Garde, A., Burkholder, D., Chi, Q., Goldstein, B., Sherwood, D.R., 2017. Identification of regulators of germ stem cell enwrapment by its niche in C. elegans. Dev. Biol. 429, 271–284. 10.1016/j.ydbio.2017.06.019.

Liu, Y., Beyer, A., Aebersold, R., 2016. On the Dependency of Cellular Protein Levels on mRNA Abundance. Cell 165, 535–550. 10.1016/j.cell.2016.03.014.

Li, X., Gordon, K.L., 2025. MIG-21 interacts with Wnt and Netrin signaling in gonad migration in C. elegans. PLoS Genet. 21, e1011866. 10.1371/journal.pgen.1011866.

Li, X., Singh, N., Miller, C., Washington, I., Sosseh, B., Gordon, K.L., 2022. The C. elegans gonadal sheath Sh1 cells extend asymmetrically over a differentiating germ cell population in the proliferative zone. eLife 11. 10.7554/eLife.75497.

Maiani, E., Milletti, G., Nazio, F., Holdgaard, S.G., Bartkova, J., Rizza, S., Cianfanelli, V., Lorente, M., Simoneschi, D., Di Marco, M., D’Acunzo, P., Di Leo, L., Rasmussen, R., Montagna, C., Raciti, M., De Stefanis, C., Gabicagogeascoa, E., Rona, G., Salvador, N., Pupo, E., Cecconi, F., 2021. AMBRA1 regulates cyclin D to guard S-phase entry and genomic integrity. Nature 592, 799–803. 10.1038/s41586-021-03422-5.

McGehee, A., 2019. The GLR-1 phenotypes of the daf-7(e1372) allele are not temperature sensitive. MicroPubl. Biol. 2019. 10.17912/micropub.biology.000158.

McIntyre, D.C., Nance, J., 2023. Niche cells regulate primordial germ cell quiescence in response to basement membrane signaling. Development 150. 10.1242/dev.201640.

McShane, E., Sin, C., Zauber, H., Wells, J.N., Donnelly, N., Wang, X., Hou, J., Chen, W., Storchova, Z., Marsh, J.A., Valleriani, A., Selbach, M., 2016. Kinetic Analysis of Protein Stability Reveals Age-Dependent Degradation. Cell 167, 803–815.e21. 10.1016/j.cell.2016.09.015.

Medwig-Kinney, T.N., Sirota, S.S., Gibney, T.V., Pani, A.M., Matus, D.Q., 2022. An in vivo toolkit to visualize endogenous LAG-2/Delta and LIN-12/Notch signaling in C. elegans. MicroPubl. Biol. 2022. 10.17912/micropub.biology.000602.

Morao, A.K., Ercan, S., 2021. Hatched and starved: Two chromatin compaction mechanisms join forces to silence germ cell genome. J. Cell Biol. 220. 10.1083/jcb.202107026.

Murphy, C.T., Hu, P.J., 2013. Insulin/insulin-like growth factor signaling in C. elegans. WormBook 1–43. 10.1895/wormbook.1.164.1.

Narbonne, P., Roy, R., 2006. Inhibition of germline proliferation during C. elegans dauer development requires PTEN, LKB1 and AMPK signalling. Development 133, 611–619. 10.1242/dev.02232.

Ouellet, J., Li, S., Roy, R., 2008. Notch signalling is required for both dauer maintenance and recovery in C. elegans. Development 135, 2583–2592. 10.1242/dev.012435.

Ow, M.C., Borziak, K., Nichitean, A.M., Dorus, S., Hall, S.E., 2018. Early experiences mediate distinct adult gene expression and reproductive programs in Caenorhabditis elegans. PLoS Genet. 14, e1007219. 10.1371/journal.pgen.1007219.

Ow, M.C., Hall, S.E., 2015. A Method for Obtaining Large Populations of Synchronized Caenorhabditis elegans Dauer Larvae. Methods Mol. Biol. 1327, 209–219. 10.1007/978-1-4939-2842-2_15.

Ow, M.C., Nichitean, A.M., Hall, S.E., 2021. Somatic aging pathways regulate reproductive plasticity in Caenorhabditis elegans. eLife 10. 10.7554/eLife.61459.

Park, D., Estevez, A., Riddle, D.L., 2010. Antagonistic Smad transcription factors control the dauer/non-dauer switch in C. elegans. Development 137, 477–485. 10.1242/dev.043752.

Park, J., Oh, H., Kim, D.-Y., Cheon, Y., Park, Y.-J., Hwang, H., Neal, S.J., Dar, A.R., Butcher, R.A., Sengupta, P., Kim, D.-W., Kim, K., 2021. CREB mediates the C. elegans dauer polyphenism through direct and cell-autonomous regulation of TGF-β expression. PLoS Genet. 17, e1009678. 10.1371/journal.pgen.1009678.

Pekar, O., Ow, M.C., Hui, K.Y., Noyes, M.B., Hall, S.E., Hubbard, E.J.A., 2017. Linking the environment, DAF-7/TGFβ signaling and LAG-2/DSL ligand expression in the germline stem cell niche. Development 144, 2896–2906. 10.1242/dev.147660.

Pierce, S.B., Costa, M., Wisotzkey, R., Devadhar, S., Homburger, S.A., Buchman, A.R., Ferguson, K.C., Heller, J., Platt, D.M., Pasquinelli, A.A., Liu, L.X., Doberstein, S.K., Ruvkun, G., 2001. Regulation of DAF-2 receptor signaling by human insulin and ins-1, a member of the unusually large and diverse C. elegans insulin gene family. Genes Dev. 15, 672–686. 10.1101/gad.867301.

Renfree, M.B., Fenelon, J.C., 2017. The enigma of embryonic diapause. Development 144, 3199–3210. 10.1242/dev.148213.

Ren, P., Lim, C.S., Johnsen, R., Albert, P.S., Pilgrim, D., Riddle, D.L., 1996. Control of C. elegans larval development by neuronal expression of a TGF-beta homolog. Science 274, 1389–1391. 10.1126/science.274.5291.1389.

Roy, D., Michaelson, D., Hochman, T., Santella, A., Bao, Z., Goldberg, J.D., Hubbard, E.J.A., 2016. Cell cycle features of C. elegans germline stem/progenitor cells vary temporally and spatially. Dev. Biol. 409, 261–271. 10.1016/j.ydbio.2015.10.031.

Schindler, A.J., Baugh, L.R., Sherwood, D.R., 2014. Identification of late larval stage developmental checkpoints in Caenorhabditis elegans regulated by insulin/IGF and steroid hormone signaling pathways. PLoS Genet. 10, e1004426. 10.1371/journal.pgen.1004426.

Seidel, H.S., Kimble, J., 2015. Cell-cycle quiescence maintains Caenorhabditis elegans germline stem cells independent of GLP-1/Notch. eLife 4. 10.7554/eLife.10832.

Seidel, H.S., Kimble, J., 2011. The oogenic germline starvation response in C. elegans. PLoS ONE 6, e28074. 10.1371/journal.pone.0028074.

Shaffer, J.M., Greenwald, I., 2022. Floxed exon (Flexon): A flexibly positioned stop cassette for recombinase-mediated conditional gene expression. Proc Natl Acad Sci USA 119. 10.1073/pnas.2117451119.

Shaner, N.C., Campbell, R.E., Steinbach, P.A., Giepmans, B.N.G., Palmer, A.E., Tsien, R.Y., 2004. Improved monomeric red, orange and yellow fluorescent proteins derived from Discosoma sp. red fluorescent protein. Nat. Biotechnol. 22, 1567–1572. 10.1038/nbt1037.

Shaner, N.C., Lambert, G.G., Chammas, A., Ni, Y., Cranfill, P.J., Baird, M.A., Sell, B.R., Allen, J.R., Day, R.N., Israelsson, M., Davidson, M.W., Wang, J., 2013. A bright monomeric green fluorescent protein derived from Branchiostoma lanceolatum. Nat. Methods 10, 407–409. 10.1038/nmeth.2413.

Shaw, W.M., Luo, S., Landis, J., Ashraf, J., Murphy, C.T., 2007. The C. elegans TGF-beta Dauer pathway regulates longevity via insulin signaling. Curr. Biol. 17, 1635–1645. 10.1016/j.cub.2007.08.058.

Simoneschi, D., Rona, G., Zhou, N., Jeong, Y.T., Jiang, S., Milletti, G., Arbini, A.A., O’Sullivan, A., Wang, A.A., Nithikasem, S., Keegan, S., Siu, Y., Cianfanelli, V., Maiani, E., Nazio, F., Cecconi, F., Boccalatte, F., Fenyö, D., Jones, D.R., Busino, L., Pagano, M., 2021. CRL4AMBRA1 is a master regulator of D-type cyclins. Nature 592, 789–793. 10.1038/s41586-021-03445-y.

Singh, N., Scott, K., Proctor, J., Gordon, K.L., 2025. An RNAi screen of Rab GTPase genes in Caenorhabditis elegans reveals that morphogenesis has a higher demand than stem cell niche maintenance for rab-1 in the somatic cells of the reproductive system. G3 (Bethesda) 15. 10.1093/g3journal/jkaf085.

Singh, N., Zhang, P., Li, K.J., Gordon, K.L., 2024. The Rac pathway prevents cell fragmentation in a nonprotrusively migrating leader cell during C. elegans gonad organogenesis. Curr. Biol. 34, 2387–2402.e5. 10.1016/j.cub.2024.04.073.

Sipe, C.W., Siegrist, S.E., 2017. Eyeless uncouples mushroom body neuroblast proliferation from dietary amino acids in Drosophila. eLife 6. 10.7554/eLife.26343.

Sood, C., Justis, V.T., Doyle, S.E., Siegrist, S.E., 2022. Notch signaling regulates neural stem cell quiescence entry and exit in Drosophila. Development 149. 10.1242/dev.200275.

Sternberg, P.W., Van Auken, K., Wang, Q., Wright, A., Yook, K., Zarowiecki, M., Arnaboldi, V., Becerra, A., Brown, S., Cain, S., Chan, J., Chen, W.J., Cho, J., Davis, P., Diamantakis, S., Dyer, S., Grigoriadis, D., Grove, C.A., Harris, T., Howe, K., Stein, L., 2024. WormBase 2024: status and transitioning to Alliance infrastructure. Genetics 227. 10.1093/genetics/iyae050.

Stewart, D., Killeen, E., Naquin, R., Alam, S., Alam, J., 2003. Degradation of transcription factor Nrf2 via the ubiquitin-proteasome pathway and stabilization by cadmium. J. Biol. Chem. 278, 2396–2402. 10.1074/jbc.M209195200.

Stiernagle, T., 2006. Maintenance of C. elegans. WormBook 1–11. 10.1895/wormbook.1.101.1.

Suarez Rodriguez, F., Sanlidag, S., Sahlgren, C., 2023. Mechanical regulation of the Notch signaling pathway. Curr. Opin. Cell Biol. 85, 102244. 10.1016/j.ceb.2023.102244.

Sulston, J.E., Horvitz, H.R., 1977. Post-embryonic cell lineages of the nematode, Caenorhabditis elegans. Dev. Biol. 56, 110–156. 10.1016/0012-1606(77)90158-0.

Sulston, J.E., Schierenberg, E., White, J.G., Thomson, J.N., 1983. The embryonic cell lineage of the nematode Caenorhabditis elegans. Dev. Biol. 100, 64–119. 10.1016/0012-1606(83)90201-4.

Taylor, S.E., 2010. Mechanisms linking early life stress to adult health outcomes. Proc Natl Acad Sci USA 107, 8507–8512. 10.1073/pnas.1003890107.

Tenen, C.C., Greenwald, I., 2019. Cell Non-autonomous Function of daf-18/PTEN in the Somatic Gonad Coordinates Somatic Gonad and Germline Development in C. elegans Dauer Larvae. Curr. Biol. 29, 1064–1072.e8. 10.1016/j.cub.2019.01.076.

Thomas, J.H., Birnby, D.A., Vowels, J.J., 1993. Evidence for parallel processing of sensory information controlling dauer formation in Caenorhabditis elegans. Genetics 134, 1105–1117.

Tian, X., Hansen, D., Schedl, T., Skeath, J.B., 2004. Epsin potentiates Notch pathway activity in Drosophila and C. elegans. Development 131, 5807–5815. 10.1242/dev.01459.

Trent, C., Tsuing, N., Horvitz, H.R., 1983. Egg-laying defective mutants of the nematode Caenorhabditis elegans. Genetics 104, 619–647. 10.1093/genetics/104.4.619.

Tullet, J.M.A., Hertweck, M., An, J.H., Baker, J., Hwang, J.Y., Liu, S., Oliveira, R.P., Baumeister, R., Blackwell, T.K., 2008. Direct inhibition of the longevity-promoting factor SKN-1 by insulin-like signaling in C. elegans. Cell 132, 1025–1038. 10.1016/j.cell.2008.01.030.

Vowels, J.J., Thomas, J.H., 1992. Genetic analysis of chemosensory control of dauer formation in Caenorhabditis elegans. Genetics 130, 105–123. 10.1093/genetics/130.1.105.

Wadsworth, W.G., Riddle, D.L., 1989. Developmental regulation of energy metabolism in Caenorhabditis elegans. Dev. Biol. 132, 167–173. 10.1016/0012-1606(89)90214-5.

Webster, A.K., Chitrakar, R., Taylor, S.M., Baugh, L.R., 2022. Alternative somatic and germline gene-regulatory strategies during starvation-induced developmental arrest. Cell Rep. 41, 111473. 10.1016/j.celrep.2022.111473.

Webster, A.K., Jordan, J.M., Hibshman, J.D., Chitrakar, R., Baugh, L.R., 2018. Transgenerational Effects of Extended Dauer Diapause on Starvation Survival and Gene Expression Plasticity in Caenorhabditis elegans. Genetics 210, 263–274. 10.1534/genetics.118.301250.

Weidemann, A., Johnson, R.S., 2008. Biology of HIF-1alpha. Cell Death Differ. 15, 621–627. 10.1038/cdd.2008.12.

Wilsterman, K., Ballinger, M.A., Williams, C.M., 2021. A unifying, eco-physiological framework for animal dormancy. Funct. Ecol. 35, 11–31. 10.1111/1365-2435.13718.

Wu, X., Fleming, A., Ricketts, T., Pavel, M., Virgin, H., Menzies, F.M., Rubinsztein, D.C., 2016. Autophagy regulates Notch degradation and modulates stem cell development and neurogenesis. Nat. Commun. 7, 10533. 10.1038/ncomms10533.

Yamamoto, S., Charng, W.-L., Bellen, H.J., 2010. Endocytosis and intracellular trafficking of Notch and its ligands. Curr. Top. Dev. Biol. 92, 165–200. 10.1016/S0070-2153(10)92005-X.

Yochem, J., Greenwald, I., 1989. glp-1 and lin-12, genes implicated in distinct cell-cell interactions in C. elegans, encode similar transmembrane proteins. Cell 58, 553–563. 10.1016/0092-8674(89)90436-4.

