## Supplemental Table and Figures for "Pre-dauer starvation rapidly and reversibly reduces niche proliferative signaling to the *C. elegans* germ line"

**Table S1. Statistical analyses for Fig. 1**
**Table S1A. All comparisons for Fig. 1B**
Kruskal-Wallis test statistic H=111.0, p<0.0001. Bolded comparisons are those displayed on graph in Fig. 1B.

| Dunn's multiple comparisons test | Mean rank diff. | Significant ? | Summary | Adjusted P Value |
|---|---|---|---|---|
| **Liquid culture vs. Starved plate** | **26.19** | **No** | **ns** | **0.0920** |
| Liquid culture vs. High density | -49.58 | Yes | **** | <0.0001 |
| Liquid culture vs. First formed | -95.59 | Yes | **** | <0.0001 |
| **Starved plate vs. High density** | **-75.77** | **Yes** | **** | **<0.0001** |
| **Starved plate vs. First formed** | **-121.8** | **Yes** | **** | **<0.0001** |
| **High density vs. First formed** | **-46.01** | **Yes** | ** | **0.0030** |

**Table S1B. All comparisons for Fig. 1C**
Kruskal-Wallis test statistic H=27.19, p<0.0001. Bolded comparisons are those displayed on graph in Fig. 1C.

| Dunn's multiple comparisons test | Mean rank diff. | Significant ? | Summary | Adjusted P Value |
|---|---|---|---|---|
| **Never-dauer vs. Starved plate** | **7.261** | **No** | **ns** | **>0.9999** |
| **Never-dauer vs. High-density plating** | **-26.61** | **Yes** | ** | **0.0014** |
| **Never-dauer vs. First-formed** | **-12.77** | **No** | **ns** | **0.4258** |
| **Starved plate vs. High-density plating** | **-33.87** | **Yes** | **** | **<0.0001** |
| Starved plate vs. First-formed | -20.03 | Yes | * | 0.0200 |
| High-density plating vs. First-formed | 13.84 | No | ns | 0.2201 |

**Table S1C. Kruskal-Wallis test for data shown in Figure 1F.**
Kruskal-Wallis test statistic H=175.3 p<0.0001.
Dunn's multiple comparison test corrected for the comparisons shown below (that is, within-genotype comparisons, not across genotype comparisons).
Bolded comparisons are those displayed on graph in Fig. 1F.

| Dunn's multiple comparisons test | Mean rank diff. | Significant ? | Summary | Adjusted P Value |
|---|---|---|---|---|
| ***daf-2(e1370)* starved vs. *daf-2(e1370)* constitutive** | **-125.4** | **Yes** | **** | **<0.0001** |
| *daf-2(e1370)* starved vs. *daf-2(e1370)* high-density | -84.58 | Yes | **** | <0.0001 |
| *daf-2(e1370)* starved vs. *daf-2(e1370)* first-formed | -99.90 | Yes | **** | <0.0001 |
| *daf-2(e1370)* high-density vs. *daf-2(e1370)* first-formed | -15.33 | No | ns | >0.9999 |
| ***daf-2(e1370)* high-density vs. *daf-2(e1370)* constitutive** | **-40.80** | **No** | **ns** | **0.2678** |
| ***daf-2(e1370)* first-formed vs. *daf-2(e1370)* constitutive** | **-25.47** | **No** | **ns** | **>0.9999** |
| ***daf-7(e1372)* starved vs. *daf-7(e1372)* constitutive** | **-147.8** | **Yes** | **** | **<0.0001** |
| *daf-7(e1372)* starved vs. *daf-7(e1372)* high-density | -97.06 | Yes | **** | <0.0001 |
| *daf-7(e1372)* starved vs. *daf-7(e1372)* first-formed | -143.8 | Yes | **** | <0.0001 |
| *daf-7(e1372)* high-density vs. *daf-7(e1372)* first-formed | -46.76 | No | ns | 0.4778 |
| ***daf-7(e1372)* high-density vs. *daf-7(e1372)* constitutive** | **-50.75** | **No** | **ns** | **0.2318** |
| ***daf-7(e1372)* first-formed vs. *daf-7(e1372)* constitutive** | **-3.990** | **No** | **ns** | **>0.9999** |

Welch’s t-test indicates mean germ cell numbers differ between constitutive dauers of *daf-2(e1370)* and *daf-7(e1372)* genotypes.
Welch’s corrected t=7.853, df=57.24
Two-tailed p<0.0001

**Table S1D. All comparisons for Fig. 1G**
Kruskal-Wallis test statistic=30.51, p<0.0001. Bolded comparisons are those displayed on graph in Fig. 1G.

| Dunn's multiple comparisons test | Mean rank diff. | Significant? | Summary | Adjusted P Value |
|---|---|---|---|---|
| **Never-dauer vs. Starved plate** | **31.77** | **Yes** | ** | **0.0012** |
| **Never-dauer vs. High density** | **0.6688** | **No** | **ns** | **>0.9999** |
| **Never-dauer vs. First-formed** | **-8.393** | **No** | **ns** | **>0.9999** |
| **Never-dauer vs. Constitutive** | **-2.119** | **No** | **ns** | **>0.9999** |
| Starved plate vs. High density | -31.10 | Yes | *** | 0.0003 |
| Starved plate vs. First-formed | -40.17 | Yes | **** | <0.0001 |
| Starved plate vs. Constitutive | -33.89 | Yes | *** | 0.0009 |
| High density vs. First-formed | -9.062 | No | ns | >0.9999 |
| High density vs. Constitutive | -2.788 | No | ns | >0.9999 |
| First-formed vs. Constitutive | 6.274 | No | ns | >0.9999 |

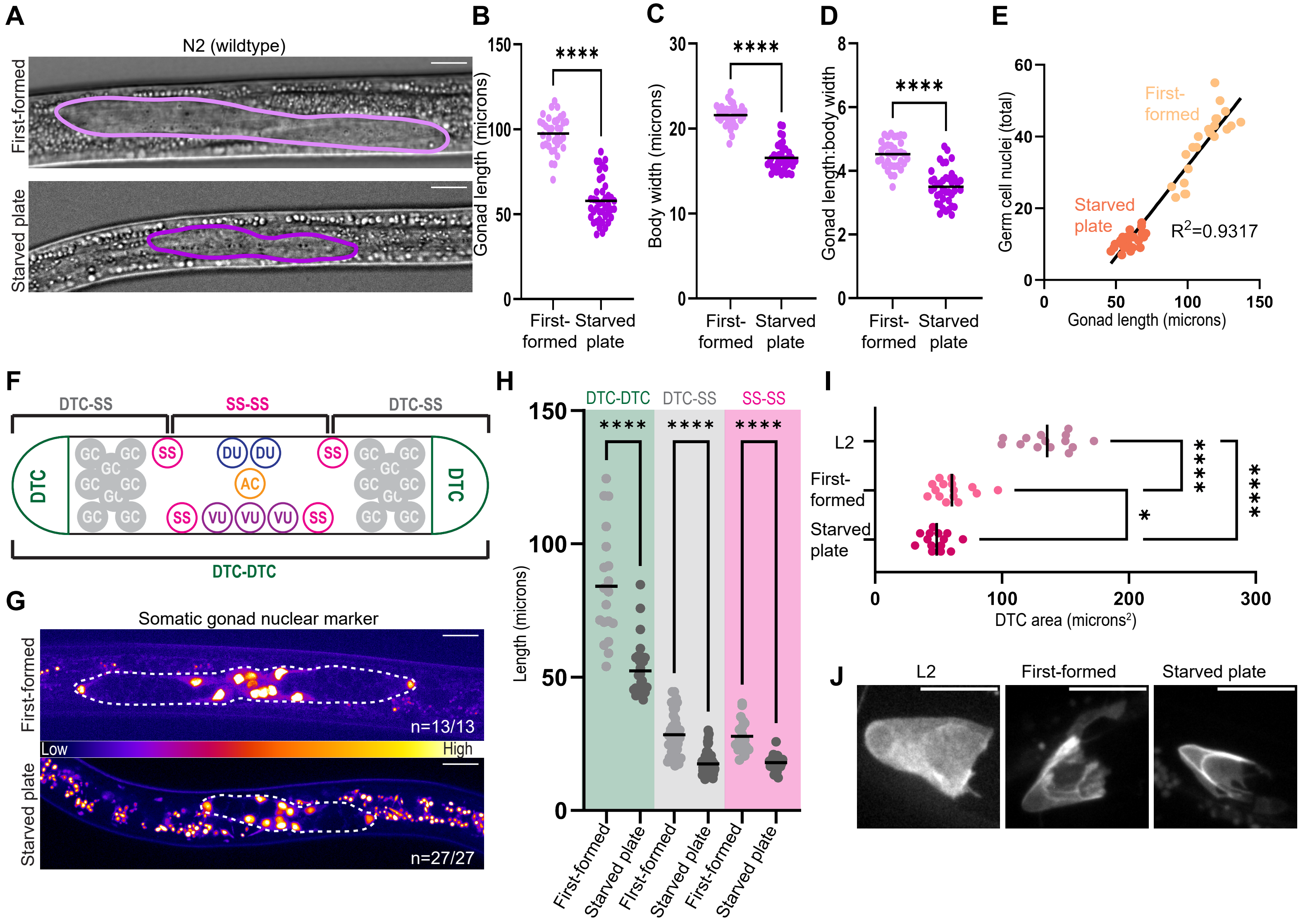


**Figure S1. Somatic gonad cell number is unaffected by method of dauer induction.**
(A) DIC images of representative wildtype N2 dauer worms also show large gonads under a treatment in which animals can feed before dauer (first-formed, top) and small gonads under a treatment in which they starve severely (starved plate, bottom). Gonads outlined in purple. This confirms that fully wildtype worms display the differential growth phenotype we observe for the marker control strain. (B) Gonad length for worms in A. (C) Body width for worms in A. (D) Ratio of gonad length to body width for worms in A showing allometric scaling between conditions. (E) Germ cell number is highly correlated to gonad length in dauer for the otherwise wild-type marker strain shown in Fig. 1 for First-formed (n=24) and starved plate (n=28) dauers, R2=0.9317. (F) Illustration of the 12 somatic gonad cells based on (Lints and Hall, 2004), L2/L3 molt). We observe the same number of cells in the dauer in G. Distal tip cells (DTC), sheath-spermathecal (SS), dorsal uterine (DU), ventral uterine (VU), anchor cell (AC). Germ cells (GC) are separated into two gonad arms. Brackets at top indicate measurements used in H. (G) Representative images of dauer gonads (outlined) with a somatic gonad-specific nuclear marker *arTi435(rps-27p::2xnls::gfp(flexon)::unc-54 3' UTR); arTi237(ckb-3p::Cre(opti)::tbb-2 3' UTR)* (Shaffer and Greenwald, 2022)). Single-slice images colored by fire LUT. First-formed (top, n=13) and starved plate treatment (bottom, n=27) all produce dauers with 12 somatic gonad cells). (F) Distance between three sets of cells of the somatic gonad in first-formed and starved dauers of the strain in (E). DTC-DTC, DTC-closest SS cell in that same arm, and SS-SS cell across the central region. DTC-DTC, Mann-Whitney test statistic U=31, p<0.0001; DTC-SS, Mann-Whitney test statistic U=193, p<0.0001; SS-SS, Mann-Whitney test statistic U=21, p<0.0001.

CONFIDENTIAL

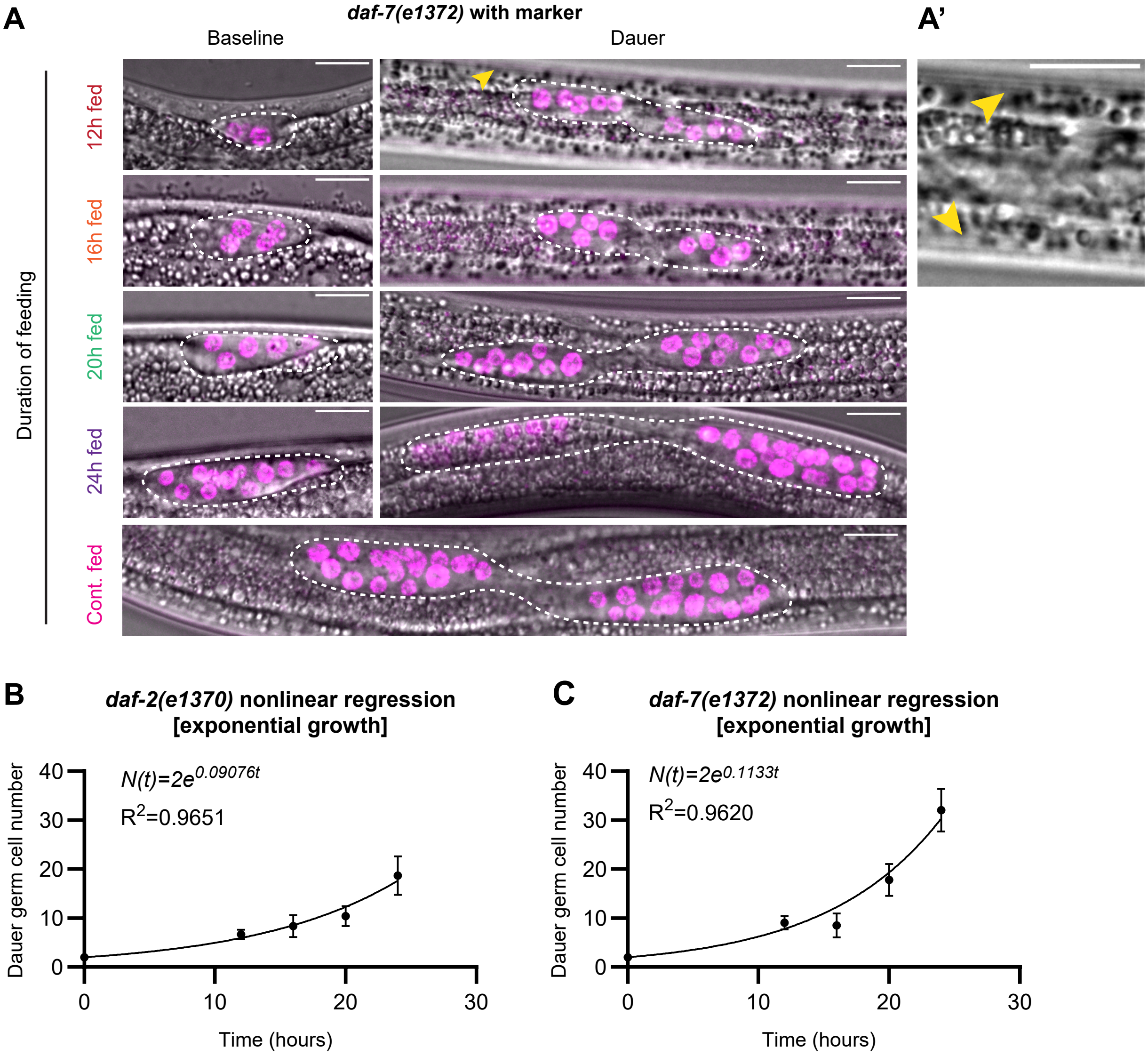


**Figure S2. *daf-7* and *daf-2* are dispensable for nutritionally-sensitive exponential gonad growth before dauer.**

(A) Representative images of *daf-7(e1372)* animals from Fig. 2C at baseline and dauer. Gonads outlined in white dashed line. Yellow arrowhead points to cuticular alae, a morphological characteristic used to verify animals were in dauer. (A') Different plane of focus of same worm shown in A for the 12h fed dauer condition, enlarged, highlighting the alae (yellow arrowheads). Animals were counted as dauer only if they had dauer morphology, including alae. (B-C) Exponential growth curves fit to average germ cell numbers in dauer shown in Fig. 2C for each number of hours fed, labeled by genotype. 0 hour datapoint has 2 primordial germ cells, a constraint of *C. elegans* gonad anatomy. Equation and R2 values displayed on plots. (B) Growth curve for *daf-2(e1370)*. (C) Growth curve for *daf-7(e1372)*.

CONFIDENTIAL

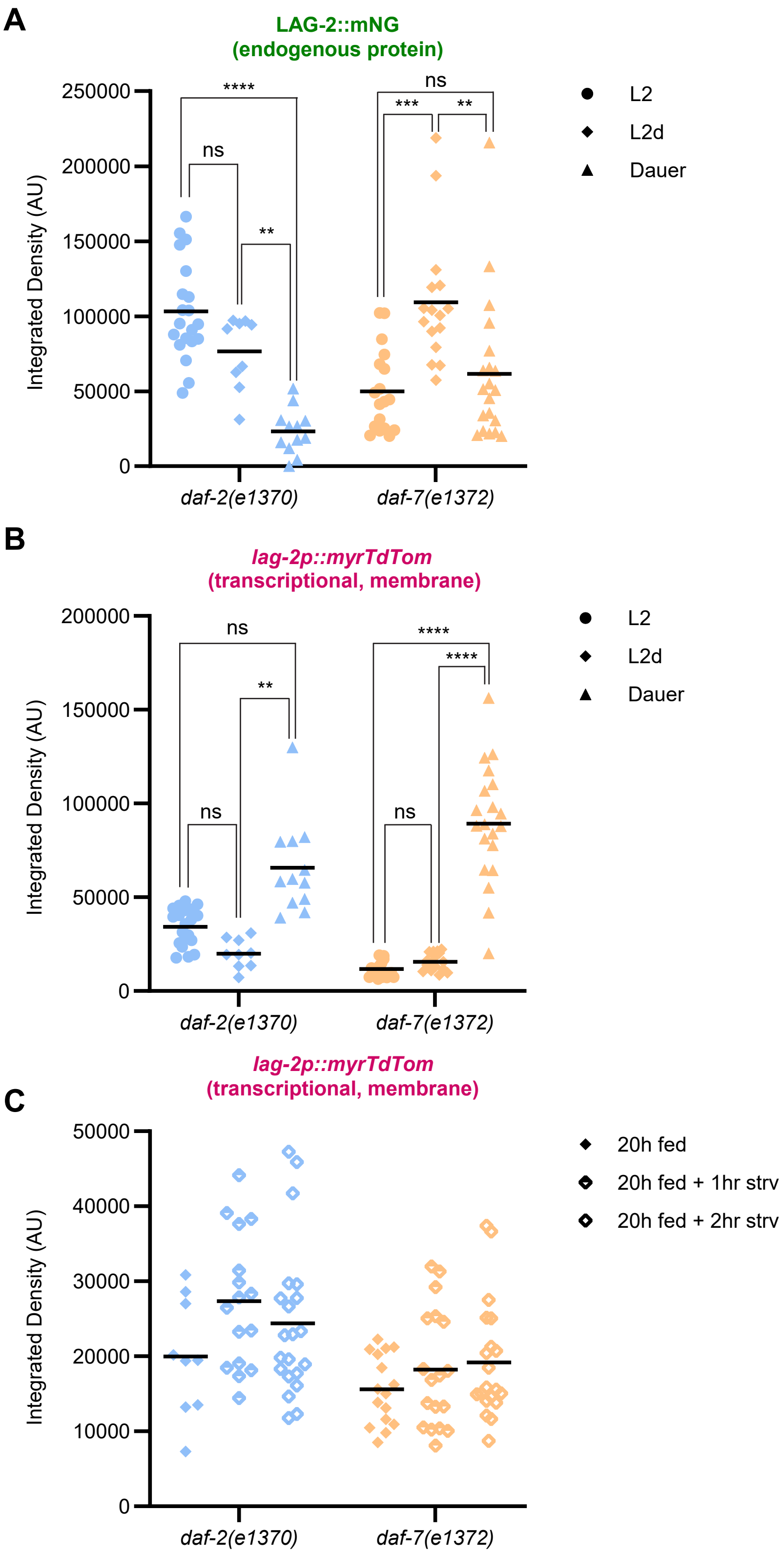


**Figure S3. Expression dynamics of *lag-2* transcriptional reporter and LAG-2::mNG protein vary across stages in *daf-2(e1370)* and *daf-7(e1372)* mutants.** (A-B) *daf-2(e1370)* (blue) and *daf-7(e1372)* (orange) mutants expressing endogenously tagged *lag-2(cp193[lag-2:: mNeonGreen])* with a membrane-localized transgenic reporter *qIs154(lag-2p:: myr::tdTomato)* at three different developmental stages, L2 larvae (reared at 16°C, as in Fig. 3H-I), L2d (reared at 25°C for 20 h, as in Fig. 3K-M, 20h fed) and in dauer (reared at 25°C until dauer). (A) *daf-2(e1370)* mutants had decreasing relative levels of LAG-2::mNG expression across the three stages, however *daf-7(e1372)* dauers has more abundant LAG-2::mNG protein in L2d than in L2 or dauer animals of that genotype. (B) Transcription of the ~3kb upstream *lag-2* promoter-driven fragment is lower in both *daf-2(e1370)* and *daf-7(e1372)* mutants at the L2 and L2d stage than in dauer. *daf-7(e1372)* dauers had higher transcriptional reporter activity (Welch's t-test t=3.189, p<0.005, mean of 89,229 A.U.) even than otherwise wildtype dauers (mean of 62,391 A.U. Fig. 3D), while *daf-2(e1370)* dauers have equivalent expression (mean of 65,776 A.U.) to wildtype dauers. This de-repression of *lag-2* in the *daf-7(e1372)* background suggests that even though low *daf-7* levels trigger dauer entry, *daf-7* acts in dauer to repress at least some regulatory elements of *lag-2* expression in the germline stem cell niche. (C) The *lag-2p::myrTdTomato* reporter does not respond to food removal on the 1- or 2-hour time scale for which LAG-2::mNG protein decays exponentially (Fig. 3M).

CONFIDENTIAL

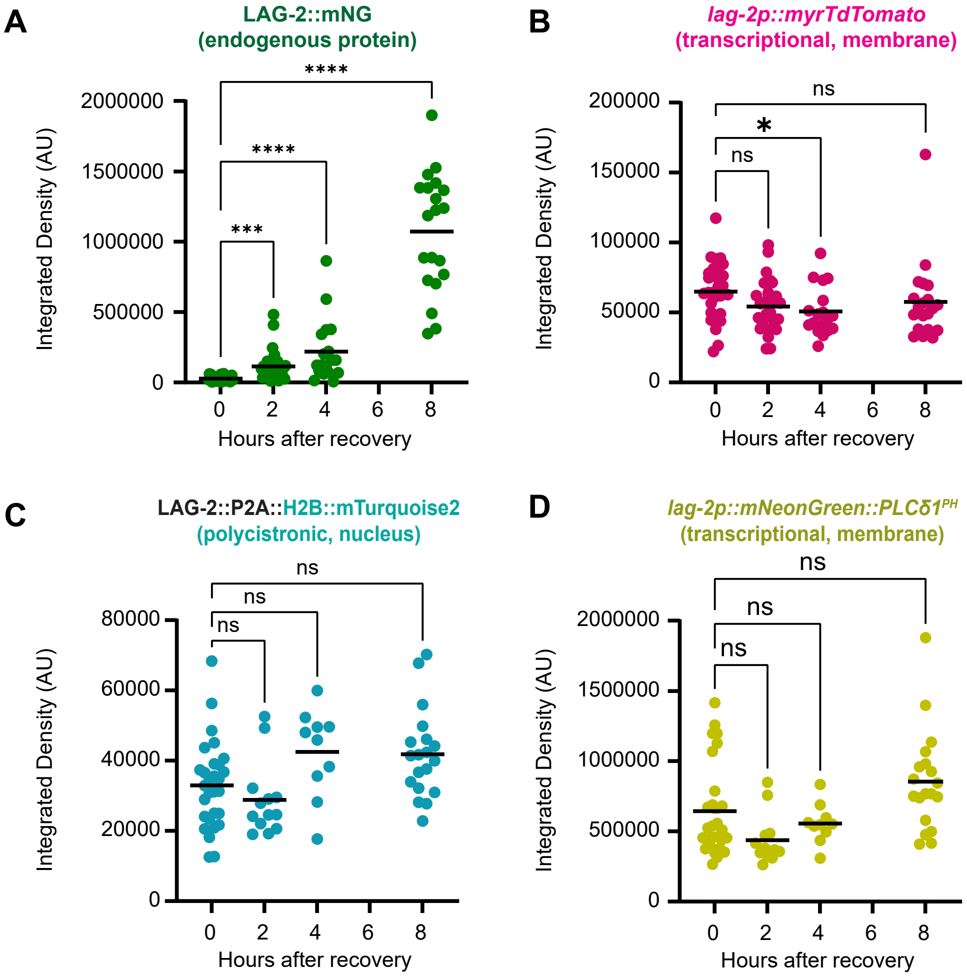


**Figure S4. Non-normalized expression of LAG-2::mNG and *lag-2* reporters during dauer recovery show no increase in transcriptional activity.**
Background-subtracted signal values in arbitrary units of fluorescence reveal a lack of absolute increase in *lag-2* transcriptional reporter activity (normalized data sets showing lack of relative increase shown in Fig. 5). (A-B) An otherwise wild-type strain coexpressing the endogenously tagged LAG-2 protein *lag-2(cp193[lag-2:: mNeonGreen])* with a membrane-localized transgenic reporter *qIs154(lag-2p:: myr::tdTomato)* recovering from dauer.
(A) LAG-2::mNG Integrated Density at 0h (0h n=30; 2h n=29; 4h=18; 8h=20). Kruskal-Wallis test statistic H=67.26, p<0.0001. Asterisks on graph indicate significance of pairwise differences by Dunn's correction for multiple comparisons as p<0.0001 ****; p<0.001 ***.
(B) Integrated density of *lag-2p::myrTdTomato* transcriptional reporter for samples in (A). Kruskal-Wallis test statistic H=8.905, p=0.0306.
(C-D) Otherwise wild-type strain co-expressing a *lag-2(bmd202 [lag-2::P2A::H2B::mTurquoise2])* polycistronic histone reporter knocked into the endogenous *lag-2* locus with a *lag-2p::mNeonGreen::PLCδ1$^{PH}$* membrane-localized transcriptional reporter.
(C) Integrated density of polycistronic reporter *lag-2(bmd202 [lag-2::P2A::H2B::mTurquoise2])* at 0h (0h n=29; 2h n=13; 4h=10; 8h=19). Kruskal-Wallis test statistic H=13.30m, p=0.0040.
(D) Integrated density of *lag-2p::mNeonGreen::PLCδ1$^{PH}$* transcriptional reporter at 0h (0h n=29; 2h n=13; 4h=10; 8h=19). Kruskal-Wallis test statistic H=17.41, p=0.0006.
For each analysis, Dunn's correction for multiple comparisons was used post-hoc to determine statistical significance of pairwise differences for relevant comparisons. For the three transcriptional reporters, no recovery timepoint was significantly increased from the 0h baseline.

CONFIDENTIAL

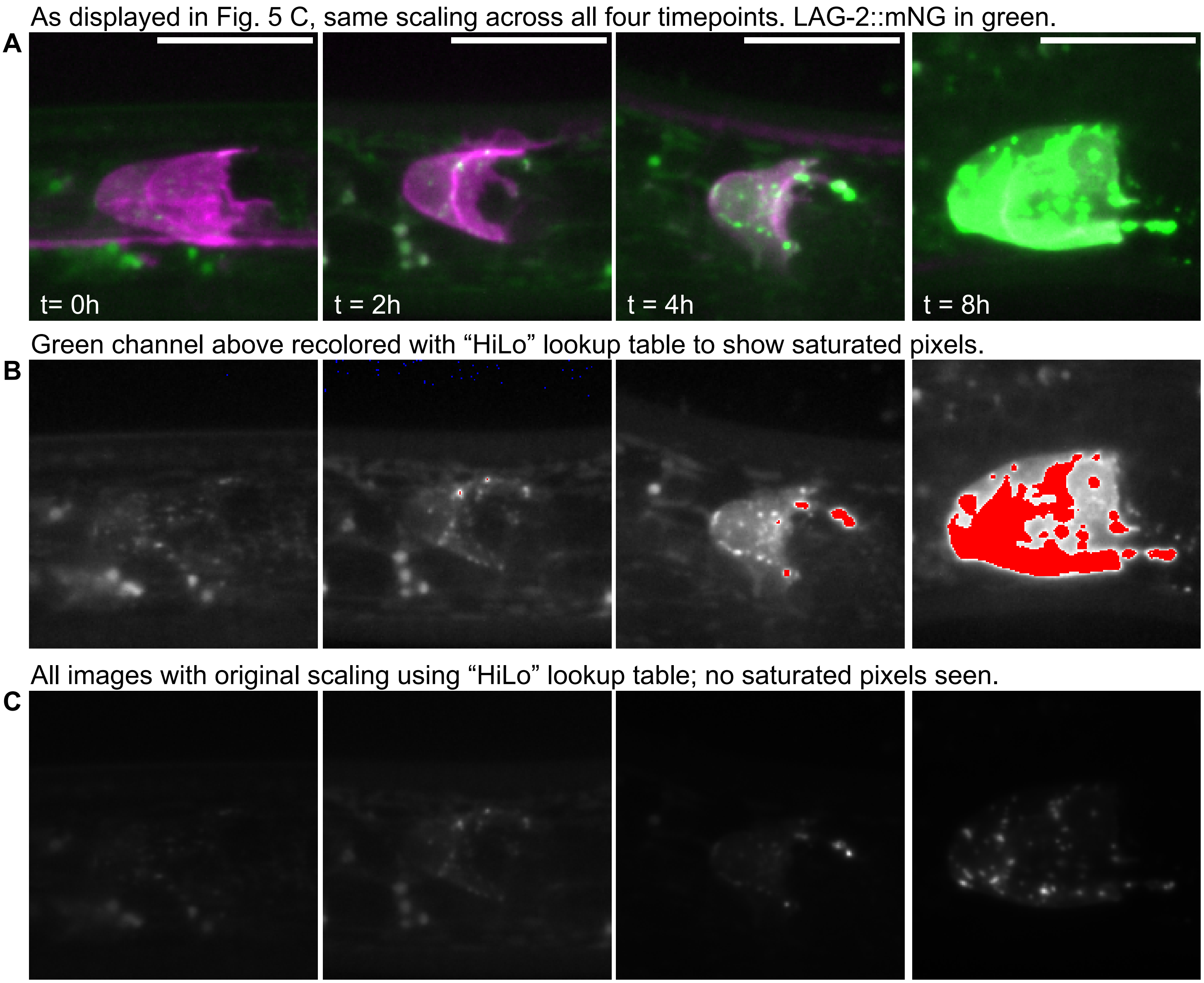


**Figure S5. Image scaling for display of qualitative comparison creates image saturation that is not present in the original data.**

An otherwise wild-type strain coexpressing the endogenously tagged LAG-2 protein *lag-2(cp193[lag-2::mNeonGreen])* with a membrane-localized transgenic reporter *qIs154(lag-2p:: myr::tdTomato)* recovering from dauer. (A) Merged images displayed in Fig. 5C. LAG-2::mNG (green), D *lag-2p:: myr::tdTomato* (magenta). Scale bar 10 μM. (B) LAG-2::mNG channel scaled identically to (A), colored with “HiLo” lookup table to show 0 value pixels (blue) and saturated pixels (red). Note saturation of brightest puncta, especially in 4h and 8h images. (C) LAG-2::mNG channel scaled individually according to the original 32-bit image scaling, also colored with “HiLo” lookup table. Note absence of saturated pixels. All fluorescence intensity measurements were made on images with original scaling before display images were produced.

CONFIDENTIAL

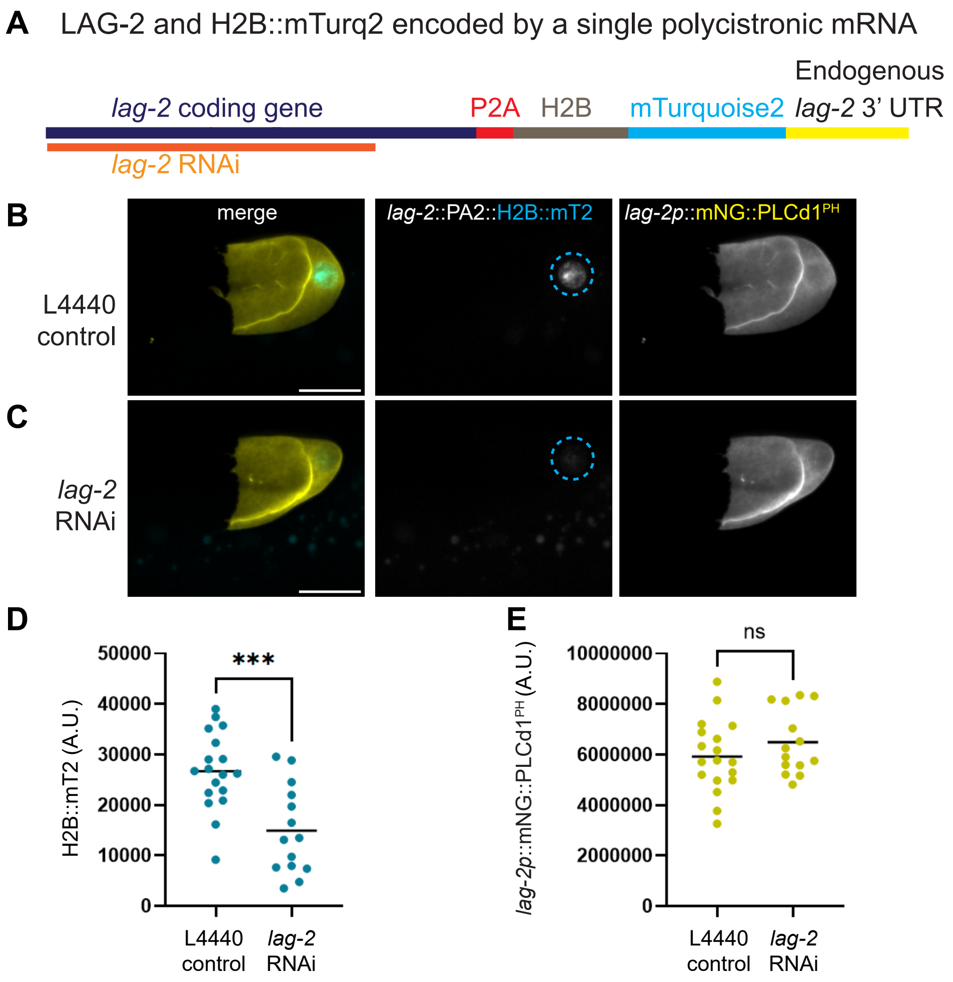


**Figure S6. Knockdown of *lag-2(bmd202 [lag-2::P2A::H2B::mTurquoise2])* signal by *lag-2* RNAi verifies H2B::mTurquoise2 transcript-based coregulation with *lag-2*.**

(A) Schematic depicting single polycistronic mRNA encoding *lag-2*, P2A "self-cleaving peptide", and H2B::mTurquoise2 targeted by *lag-2* RNAi (orange) to demonstrate the principle of coregulation of LAG-2 and H2B::mTurq2 at the transcript level. See Methods.

(B-C) Sum projection through 6 µm Z depth of DTCs of L4 worms coexpressing *lag-2p::mNeonGreen::PLCδ1$^{PH}$* (yellow in merge, right column) and *lag-2(bmd202 [lag-2::P2A::H2B::mTurquoise2])* (cyan in merge, center column) reared for 24 h from the the L2 stage on *E. coli* HT115 bacterial food expressing L4440 control vector (n=18) or *lag-2* RNAi (n=14). All images of each fluorescence channel scaled the same across treatments. All scale bars 10 µm.

(B) DTC after treatment with control vector L4440.

(C) DTC after treatment with *lag-2* RNAi.

(D-E) Background-subtracted integrated density fluorescence intensity measurements were made for the nuclei (D) and membrane (E) of one DTC per animal.

(D) H2B::mTurquoise2 signal shows ~44% knockdown after 24 hours of *lag-2* RNAi exposure. Welch's corrected t = 3.979, df=25.98, p<0.005.

(E) Transcriptional reporter membrane expression in the DTC is not affected by *lag-2* RNAi knockdown, Welch's corrected t = 1.173, df=29.35, p=0.25.

CONFIDENTIAL

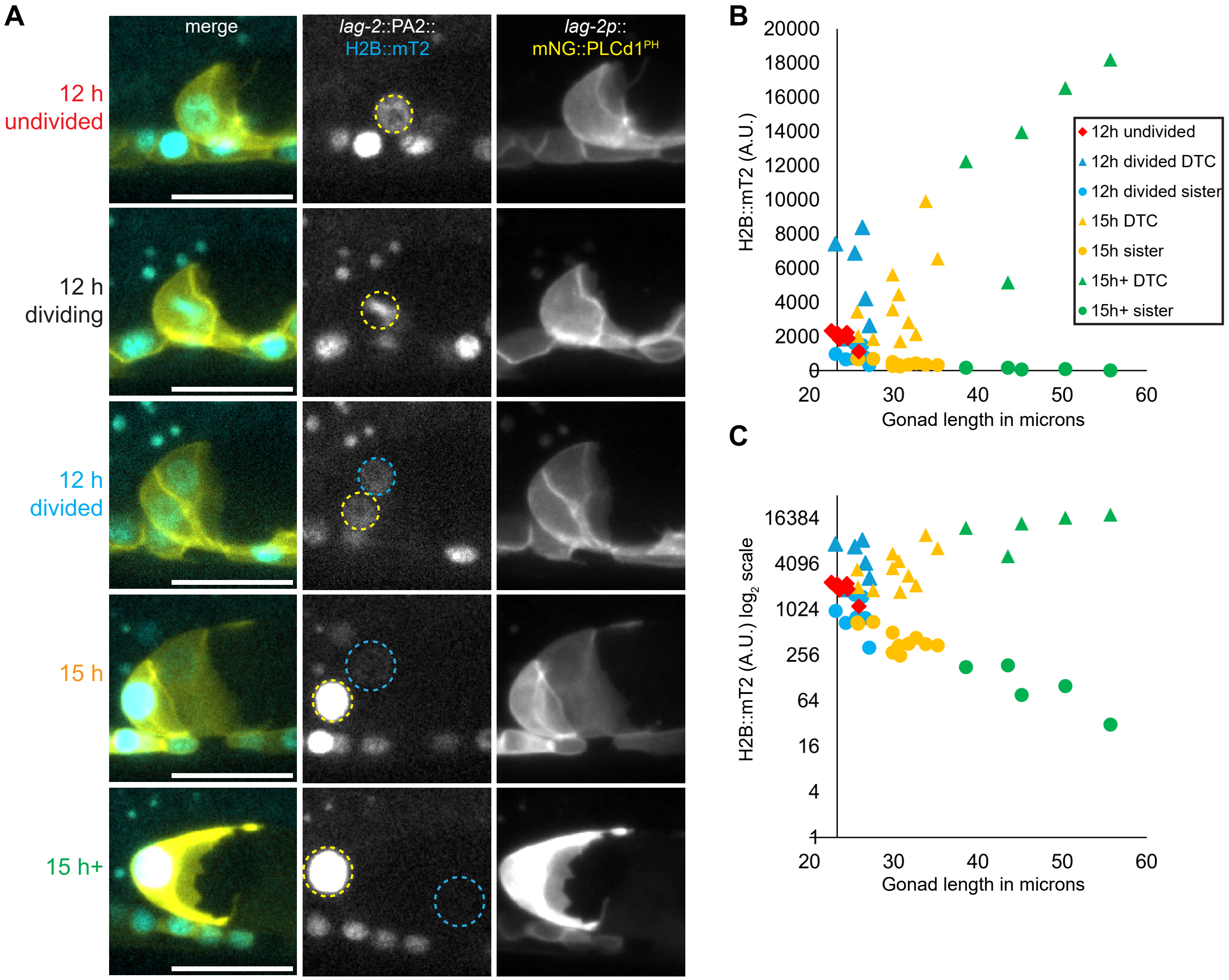


**Figure S7. Signal decay of *lag-2(bmd202 [lag-2::P2A::H2B::mTurquoise2])* reveals effective detection of continuous vs. previous expression and transcript-based coregulation with *lag-2*.**
(A) The DTC and its progenitor cells (Z1, Z1.a, Z1.aa, and Z4, Z4p, Z4.pp) express *lag-2*, while the DTC sisters (Z1.ap and Z4.pa) do not. Sum projections through ~3.3 microns of the gonad tip for worms coexpressing *lag-2p::mNeonGreen::PLCδ1*$^{PH}$ (yellow in merge, right column) and *lag-2(bmd202 [lag-2::P2A::H2B::mTurquoise2])* (cyan in merge, center column) 12 h after release from L2 arrest for which Z1.a or Z4.p had not divided (top row, yellow circles, n=6), was actively dividing (second row, yellow circle, n=1), or had just divided (third row, n=7). DTC (Z1.aa or Z4.pp, yellow circle) and its sister cell (Z1.ap or Z4.pa, blue circle) dramatically diverge in H2B::mTurquoise2 signal three hours later (15 h post release from L1 arrest, fourth row, n=11). By later L2, no fluorescence was easily visible in the SS cells (fifth row, n=5). Representative images chosen that had close-to-average gonad length and DTC signal for that stage. All images of each fluorescence channel scaled the same across stages. All scale bars 10 μM.
(B-C) Background-subtracted fluorescence intensity measurements were made for the nuclei of these cells and plotted against tip-to-tip gonad length as a proxy for developmental time. Y-axis crosses at the gonad length of the single specimen with an actively dividing Z1.a cell (shown in second row of A).
(B) Plotting these measurements on a linear scale reveals the linear increase over time of H2B::mTurquoise2 *lag-2* transcriptional signal in the DTC (triangles). Even at the earliest timepoint for which we can observe both the DTC and its sister (blue data points), the DTC is on average ~4-fold brighter than the cell in which lag-2 is no longer actively expressed, demonstrating that while H2B::mTurquoise detectably perdures after it is made, newly synthesized H2B::mTurquoise2 quickly swamps that inherited signal.
(C) Plotting these measurements on a $\log_2$ scale reveals the exponential decay of H2B::mTurquoise2 signal in the sister of the DTC (circles). The signal has a half-life of ~3h (12h average of 955 A.U., 15 h average of 447 A.U., factor of 0.46).

CONFIDENTIAL
